# HssS activation by membrane heme defines a paradigm for 2-component system signaling in *Staphylococcus aureus*

**DOI:** 10.1101/2022.11.03.515132

**Authors:** Vincent Saillant, Léo Morey, Damien Lipuma, Pierre Boëton, Pascal Arnoux, Delphine Lechardeur

**Affiliations:** Micalis Institute, INRAE, AgroParisTech, Université Paris-Saclay, 78350 Jouy-en-Josas, France; Aix Marseille Université, CEA, CNRS, BIAM, 13108, Saint Paul-Lez-Durance, France

**Author notes:** Note: in this report, heme refers to iron protoporphyrin IX regardless of the iron redox state, whereas hemin refers to ferric iron protoporphyrin IX.

**Keywords:** heme, *Staphylococcus aureus*, two-component system, virulence, membrane

## Abstract

Strict management of intracellular heme pools, which are both toxic and beneficial, can be crucial for bacterial survival during infection. The human pathogen *Staphylococcus aureus* uses a two-component heme sensing system (HssRS), which counteracts environmental heme toxicity by triggering expression of the efflux transporter HrtBA. The HssS heme sensor is a HisKA-type histidine kinase, characterized as a membrane-bound homodimer containing an extracellular sensor and a cytoplasmic conserved catalytic domain. To elucidate HssS heme sensing mechanism, a structural simulation of the HssS dimer based on Alphafold2 was docked with heme. In this model, heme is embedded in the membrane bilayer with its 2 protruding porphyrin propionates interacting with 2 conserved Arg94 and Arg163 that are located extracellularly. Mutagenesis of these arginines and of 2 highly conserved phenylalanines, Phe25 and Phe128, in the predicted hydrophobic heme binding pocket abolished the ability of HssS to induce HrtBA synthesis. This study gives evidence that exogenous heme interacts with HssS at the membrane/extracellular interface to initiate HssS activation to induce HrtBA-mediated heme extrusion from the membrane. This “gatekeeper” mechanism could limit intracellular diffusion of exogenous heme in *S. aureus*, and may serve as a paradigm for how efflux transporters control detoxification of exogenous hydrophobic stressors.

**Importance:** In the host blood, pathogenic bacteria are exposed to the red pigment heme that concentrates in their lipid membranes, generating cytotoxicity. To overcome heme toxicity, *Staphylococcus aureus* expresses a membrane sensor protein, HssS. Activation of HssS by heme triggers a phosphorelay mechanism leading to the expression of a heme efflux system, HrtBA. This detoxification system prevents intracellular accumulation of heme. Our structural and functional data reveal a heme-binding hydrophobic cavity in HssS within the TM helices at the interface with the extracellular domain. This structural pocket is important for the function of HssS as a heme sensor. Our findings provide a new basis for the elucidation of pathogen sensing mechanisms as a prerequisite to the discovery of inhibitors.

## Introduction

*Staphylococcus aureus* is a Gram-positive opportunist bacterium that asymptomatically colonizes the skin and nostrils of nearly one-third of the human population (1). However, this organism can breach defensive barriers in the compromised host to cause invasive diseases such as endocarditis, toxic shock syndrome, osteomyelitis, and sepsis (1). Like most pathogens, successful infection by *S. aureus* involves the production of numerous virulence determinants including toxins, immune-modulatory factors, and exoenzymes, and requires expression of factors that facilitate adaptation to the varied host environments (2–4).

Heme, an iron containing tetrapyrrole, is the bioactive cofactor of blood hemoglobin (Hb) (5). The importance of heme resides in the unique properties of its iron center, including the capacity to undergo electron transfer, perform acid-base reactions, and interact with various coordinating ligands (5, 6). On the other hand, redox reactions of heme iron with oxygen generate reactive oxygen species (ROS), which provoke damage to proteins, DNA and lipids (5, 6). Since heme is hydrophobic and cytotoxic, its concentration and availability must be tightly regulated.

In addition to utilizing heme as a nutrient iron source, *S. aureus* can employ both endogenously synthesized and exogenously acquired heme as a respiratory cofactor (7–9). Heme toxicity is offset in *S. aureus* by a conserved strategy for heme detoxification and homeostasis involving a heme-regulated efflux pump (HrtBA; Heme-regulated transport). HrtBA was also identified in *Enterococcus faecalis*, *Lactococcus lactis, Streptococcus agalactiae*, *Bacillus anthracis*, and *Corynebacterium diphtheriae* (10–13). HrtB topology classifies this permease as a MacB-like ABC transporter (14, 15). Rather than transporting substrates across the membrane, MacB couples cytoplasmic ATP hydrolysis with transmembrane conformational changes to extrude substrates from the periplasmic side or from the lateral side of its transmembrane domains (TM) (15, 16). A heme-binding site was recently identified in the outer leaflet of the HrtB dimer membrane domain from which heme is excreted (17).

HrtBA expression in numerous Gram positive pathogens is managed by HssRS (Hss; <u>h</u>eme <u>s</u>ensing <u>s</u>ystem), a two-component system (TCS) (13, 18–20). HssS senses heme presence in the environment and transduces the signal to HssR, the transcriptional regulator of *hrtBA* (13, 19, 20). HssS is a prototypical HisKA histidine kinase with a short Nt cytoplasmic domain, and two TM helices flanking a 133 amino acid (AA) extracellular domain (19). The Ct cytoplasmic domain is organized in structurally conserved modules: the HAMP domain (present in <u>H</u>istidine kinases, <u>A</u>denylate cyclases, <u>M</u>ethyl accepting proteins and <u>P</u>hosphatases) connects the second TM to the dimerization and histidine phosphorylation domains (HisKA). A catalytic and ATP-binding (HATPase c) domain lies at the carboxyl terminus. Upon activation, HssS undergoes autophosphorylation of the His-249 residue and subsequently transfers the phosphoryl group to the Asp-52 residue of the HssR response regulator (19). As HssS is activated by environmental heme, it is assumed that the HK extracellular domain (ECD) harbors the sensing function (21).

Here, we show that membrane heme, rather than extracellular heme, is the activating signal for the transient activation of HssS in *S. aureus*. To identify a domain within HssS that may participate in heme signal reception, we performed a structural simulation of the dimer which was docked with heme. A single conserved hydrophobic structural domain (per monomer) with 2 conserved anchoring arginines at the interface between the membrane and the extracellular domain were predicted to accommodate heme. Based on this approach, we performed targeted mutagenesis and identified pivotal residues required for HssS binding to heme. Our work reveals a new mechanism of direct ligand sensing of a histidine kinase at the membrane level. We conclude that membrane heme control of HssS combined with membrane heme extrusion by HrtB constitute a defense system for bacteria when they are exposed to lysed erythrocytes.

## Results

### *hrtBA* induction is the readout for HssS activation

Heme conditions leading to expression of *hrtBA* and *hssS* were assessed. For this, an *S. aureus* HG001 Δ*hssRS* mutant was transformed with plasmid p*hssRS-*HA, encoding HssR and a C-terminal HA-tagged version of HssS (Table S1). Antibodies against HA and HrtB were used for detection (Fig. 1A). Amounts of HrtB were increased in the presence of heme, as reported in *S. aureus* (13), while HssS expression remained constant (Fig. 1A). Accordingly, *hssRS* promoter activity, measured by β-gal expression from a P*_hssRS_*-*lac* fusion (pP*_hssRS_*-*lac*, Table S1), was independent of heme concentration (Fig. 1B). In contrast, the P*_hrtBA_* reporter (pP*_hrtBA_*-*lac;* Table S1) responded linearly with increasing concentrations of exogenously supplied hemin (Fig. 1C). As P*_hrtBA_* was specifically activated by constitutively expressed HssRS TCS (Fig. 1D), these data establish P*_hrtBA_* induction as a specific reporter of HssRS heme sensing and signaling.

**Figure 1.**
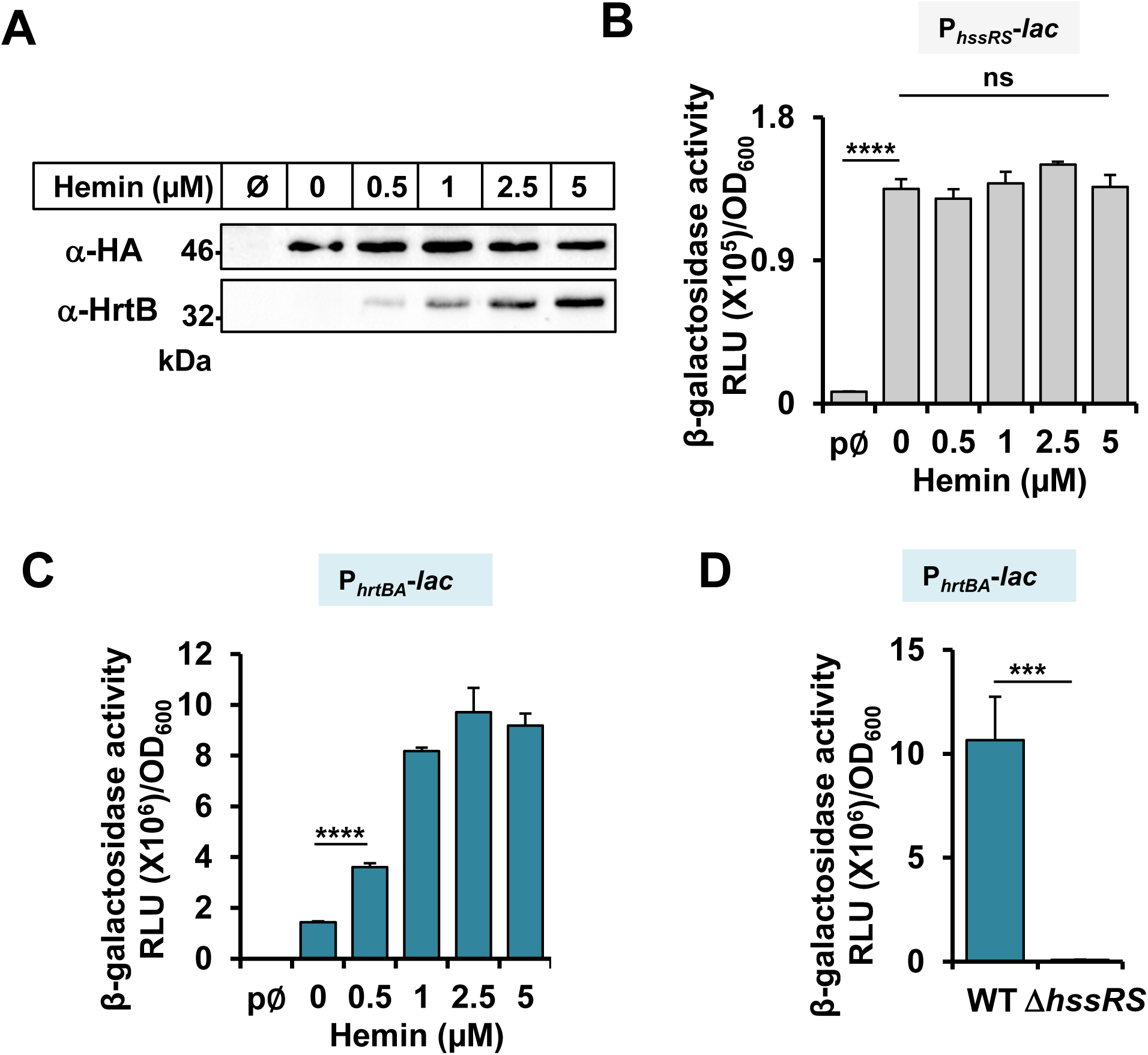
P*_hrtBA_* induction reports HssS activity. (A) HssS and HrtB expressions in presence of heme. *S. aureus* HG001 Δ*hssRS*(p*hssRS*-HA) was incubated with the indicated concentrations of hemin. WB on bacterial lysates was performed with anti-HA and anti-HrtB antibodies. Result is representative of 3 independent experiments. (B, C) *hssRS* and *hrtBA* transcription regulation by exogenous heme. *S. aureus* HG001 transformed with pP*_hssRS_*-*lac* (B) or pP*_hrtBA_*-*lac* (C) were grown in BHI to OD_600_ = 0.5 prior addition of the indicated concentration of hemin for 1.5 h. β-gal activity was quantified by luminescence. Results represent the average ± S.D. from triplicate independent experiments. ****, *P <*0.0001; ns, non-significant, Student’s *t* test. (D) P*_hrtBA_* is not induced in the HG001 Δ*hssRS* mutant. WT and Δ*hssRS* HG001 strains transformed with pP*_hrtBA_*-*lac* were grown and β-gal activity determined as in (B-C) with 1 µM hemin. Results represent the average ± S.D. from triplicate independent experiments. ***, *P <*0.001, Student’s *t* test.

### HssS transient activation by hemin signals intracellular heme accumulation

To get insights into the mechanism of heme sensing by HssS, we followed the kinetics of HssS stimulation by heme in WT HG001 with the fluorescent reporter (pP*_hrtBA_*-GFP) (Table S1). Hemin addition led to a transient P*_hrtBA_* response at the beginning of HG001 growth, with a maximal response output within a few hours post heme addition (Fig. 2A). No fluorescence was detected in the strain carrying the promoterless plasmid (data not shown). At toxic heme concentrations, P*_hrtBA_* induction kinetics seems to follow the growth delay and reaches highest induction in the presence of 1 and 2.5 µM heme (Fig. 2A). P*_hrtBA_* induction phase was followed by a marked drop in fluorescence, likely corresponding to termination of the P*_hrtBA_* induction phase (Fig. 2A). In stationary phase bacteria, fluorescence associated to pP*_hrtBA_*-GFP expression stabilized following induction by heme, confirming the transient activation by HssS (Fig. S1A). Expression kinetics of GFP and HrtB correlated as shown on WB (Fig. S1B).

**Figure 2.**
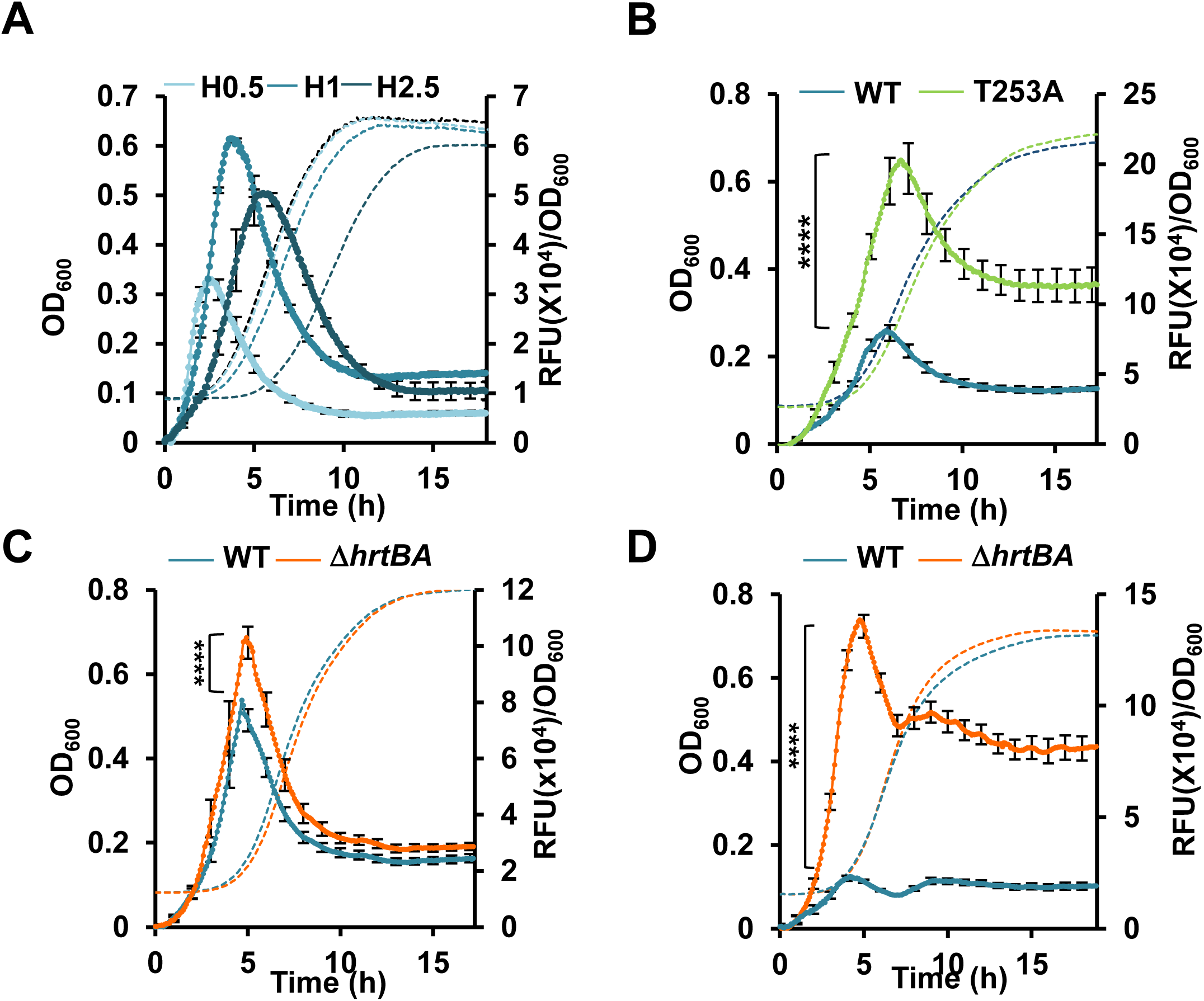
HssS transient activation reports intracellular accumulation of exogenous heme. (A) Dynamics of P*_hrtBA_* induction in *S. aureus* HG001 by exogenous heme. Bacteria transformed with pP*_hrtBA_*-GFP were diluted from an ON culture to OD_600_ = 0.01 in CDM with the indicated concentration of hemin in a microplate spectrofluorimeter Infinite (Tecan). Both fluorescence (Exc: 475 nm; Em: 520 nm) and OD_600_ were recorded every 5 min for the indicated time. Fluorescence (RFU) at each time point was divided by the corresponding OD_600_. Results of hemin induced fluorescence minus non-induced (background, 0 µM hemin) are displayed. Results represent the average ± S.D. from triplicate biological samples. The corresponding growth curves are shown. (B) Phosphatase activity of HssS. Δ*hssRS* HG001 transformed with pGFP(HssS) or pGFP(HssS T253A) were diluted in CDM ± 1 µM hemin from an ON culture in 96-microplate as in (A). Fluorescence emission was quantified as in (A). Fluorescence (RFU) at each time point was divided by the corresponding OD_600_. Results of hemin induced fluorescence minus non-induced (background, 0 µM hemin) are displayed. Results represent the average ± S.D. from triplicate biological samples. ****, *P <* 0.0001, Student’s *t* test. The corresponding growth curves are shown. (C) Dynamics of P*_hrtBA_* induction in WT and Δ*hrtBA* HG001 strains by exogenous heme. *S. aureus* HG001 WT and Δ*hrtBA* strains transformed with pP*_hrtBA_*-GFP were diluted from an ON culture to OD_600_ = 0.01 in CDM with 1 µM hemin in a 96 well microplate. OD_600_ and GFP expression were followed in a spectrofluorimeter Infinite (Tecan) as in Fig. 1. Results of hemin induced fluorescence minus non-induced (background, 0 µM hemin) are displayed Results represent the average ± S.D. from triplicate biological samples. The corresponding growth curve are shown. ****, *P <* 0.0001, Student’s *t* test. (D) Dynamics of HssS activation in HG001 WT and Δ*hrtBA* strains by hemoglobin. Fluorescence emission kinetic was followed as in (C) in HG001 WT and Δ*hrtBA* strains transformed pP*_hrtBA_*-GFP with 0.25 µM human hemoglobin (equivalent to 1 µM hemin) added to the culture medium. Results represent the average ± S.D. from triplicate technical samples and are representative of 3 independent experiments. ****, *P <*0.0001, Student’s *t* test. The corresponding growth curves are shown.

Negative control of HisKA is provided by its phosphatase activity on the phosphorylated regulator. The conserved catalytic histidine residue together with the adjacent conserved threonine residue (HXXXT motif) is a key target for HisKA phosphatases (22, 23). To evaluate the role of HssS phosphatase activity in its activation dynamics, the threonine T253 was mutated to alanine in the plasmid p*hssRS*-HA, P*_hrtBA_*-*gfp* (pGFP(HssS)) to generate pGFP(HssS T253A) (Table S1). HG001 Δ*hssRS* was then transformed with both plasmids and response to heme was characterized. Higher fluorescence emission was observed in the strain expressing the HssS T253A allele compared to the WT strain, but remained transient (Fig. 2B). This result illustrates the duality of HssS as a kinase and phosphatase at any time point in response to its activation by heme.

To test the possibility that transient HssS activation was related to HrtBA-mediated heme efflux, kinetics of GFP expression from P*_hrtBA_* in Δ*hrtBA* and WT strains by subtoxic heme concentrations were compared (Fig. 2C). GFP expression was also transient in the Δ*hrtBA* strain, indicating that HrtBA expression did not explain transient HssS activation. While P*_hrtBA_* dynamics were similar in both strains, fluorescence emission reached higher values in Δ*hrtBA* (Fig. 2C). We hypothesize that HssS activation intensity is correlated to the intracellular accumulation of heme upon *hrtBA* deletion. To test this, heme accumulation was visualized by the color of culture pellets from the Δ*hrtBA* mutant compared to the WT strain exposed to hemin (Fig. S2A). Accordingly, the Δ*hrtBA* mutant accumulated about twice more heme than did the WT as evaluated by the pyridine hemochrome assay (Fig. S2B). These results show that intracellular heme pools impact HssS activation, raising the question of where the heme-HssS interface is localized.

Replacing free hemin by Hb led to a fluorescence emission intensity that was more than 5 times higher in the Δ*hrtBA* strain than in the WT strain (Fig. 2D). Interestingly, the kinetics of GFP expression were modified in the presence of Hb compared to hemin. Slow and continuous delivery of hemin from Hb compared to a fast and short delivery of free hemin to the bacteria could provide an explanation to the observed distinct kinetics of GFP expression and transient HssS activation. These results further correlate membrane HssS activation with intracellular heme accumulation.

The documented role of HssS as the signal transmitter required for HrtBA expression gives strong *in vivo* evidence that HssS activation involves the pool of *S. aureus*-associated heme rather than exclusively extracellular heme as generally considered (18–20). However, as exogenous heme is detectable extracellularly, in the membrane, and in the cytoplasm (10, 24, 25), we cannot discriminate which bacterial compartment drives HssS activation.

### Heme docking on a prediction model of HssS reveals a putative heme binding region at the interface between membrane and extracellular domains

Attempts to identify specific AAs residues within HssS that may participate in heme signal reception have been hampered by the lack of an experimental three-dimensional structure. We relied on an *in silico* approach using HssS structure prediction by AlphaFold2 (AF2). Results of AF2 inferencing generated a model with a mean predicted local distance difference test (pLDDT) value of 88 (https://alphafold.ebi.ac.uk/entry/A5IVE3). This indicates a confident prediction, according to guidelines set out on EMBL’s AlphaFold Protein Structure Database, available at https://alphafold.ebi.ac.uk. Dimer prediction was obtained with Alphafold advanced (https://colab.research.google.com/github/sokrypton/ColabFold/blob/main/beta/AlphaFold2_advanced.ipynb) (Fig. 3A). The structural cytoplasmic domains of the AF2 model were consistent with previously assigned domain predictions (based on InterProScan annotations) as displayed by a prototypical HisKA (Fig 3A). The overall structure of HssS ECD is a mixed α/ β fold with a PDC (PhoQ-DcuS-CitA)-like structure topology (26) (Fig. 3B). The central 4-stranded antiparallel β-sheet is flanked by α−helices on either side; a long N-terminal α-helix and a short C-terminal α-helix that both lie on the same side of the sheet (Fig. 3B). The long N-terminal helix is initiated by TM helix α1 (identified as residues [11-31] by Orientation of Proteins in Membranes (OPM, https://opm.phar.umich.edu/) and then participates in the mixed α/β fold of the PDC domain. A second TM helix (helix α4) (identified as residues [166-187]) allows the polypeptide chain to run in the intracellular space toward the HAMP and the HisKA domains (Fig. 3B).

**Figure 3.**
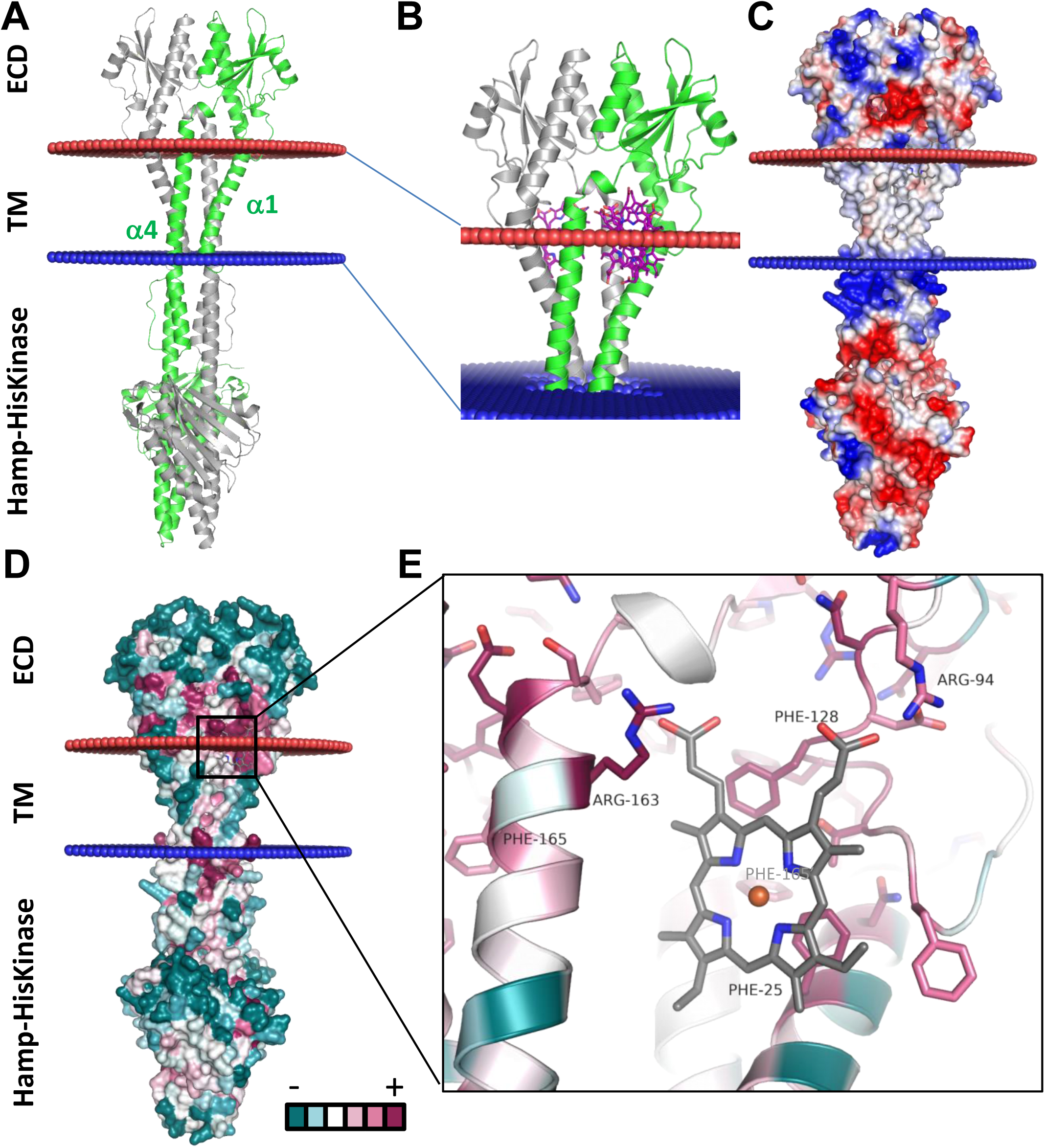
Alphafold structural model the HssS and heme docking solutions. (A) Alphafold model of *Staphylococcus aureus* HssS dimer, with one chain colored in green and the other in grey. The position of the membrane proposed by the OPM server is shown by red and blue spheres. (B) Superimposition of all the docking solutions using the ECD domains of HssS. These solutions are all < −8 kcal/mol and suggest the presence of two docking sites, which represents a single binding site due to the two-fold symmetry of the dimer. This binding site is located at the interface between the membrane and the extracellular space. (C) Electrostatic surface potential calculated with APBS with a ramp from −5 kTe (red) to +5 kTe (blue). (D) Mapping of sequence conservation on the surface of the model depicts only a few solvent exposed conserved residues. (E) Superimposition of the best docking solution of heme (colored in grey) with with predicted residues within 5Å.Cartoon and residues are colored according to sequence conservation (Fig. 3A) and only conserved residues are shown in stick.

We then used the program AutoDock Vina (https://ccsb.scripps.edu/) to dock heme on the surface of the HssS dimer. Docking used either the intracellular domains or the membrane and extracellular domains (Fig. 3B and Fig. S3). Using the intracellular part of HssS, all the docking solutions are above −8 kcal/mol and are scattered on the surface of the protein (Fig. S3). On the contrary, using the ECDs, all the docked solutions are below −8 kcal/mol and fall on two areas that are related by the two-fold symmetry of the dimer, therefore representing a single binding site (Fig. 3B). This binding site located at the interface between the lipid bilayer and the extracellular space and defined by 2 helices α1 and α4, together with an internal loop within the ECD (Fig. 3B-D). This predicted heme binding site is apolar on most of its surface, except for the top that is lined with positive and negative charges (Fig. 3D and Fig. 3E). In the best docking solution (−10.1 kcal/mol), the two heme propionates would be able to engage in a salt-bridge, one with the conserved Arg94, the other with the conserved Arg163 (Fig. 3E and Fig. S4). Furthermore, heme is surrounded by a few highly conserved hydrophobic residues (Phe25, Phe128, and Phe165 belonging to the second monomer (Phe’165) being below 4Å from the porphyrin ring (Fig. 3E and Fig. S4). This position did not reveal the usual AAs that coordinate heme such as histidine, methionine or tyrosine. Docking of heme on *S. epidermidis* HssS structural AF2 prediction (which shares 64 % identity with *S. aureus* HssS) identified the same binding position (data not shown).

We next used an *hssS* mutational approach to challenge the proposed model of HssS heme recognition.

### Conservation of the predicted heme binding domain is determinant for HssS activation

We first examined the importance of the 2 conserved arginines Arg94 and Arg163 in heme docking to HssS by generating alanine substitutions in pGFP(HssS) (Table S1). The three constructs pGFP(HssS), pGFP(HssS R94A) and pGFP(HssS R163A) (Table S1) were established in the HG001 Δ*hssRS* mutant. Activation by heme of either HssS R94A or R163A was strongly diminished compared to the WT histidine kinase as shown by the diminished fluorescence kinetic response (Fig. 4A). While expression levels of HssS R94A, HssS R163A, and WT HssS were similar, expression of HrtB was strongly decreased in HssS point mutants (Fig. 4B). Accordingly, the 2 arginine HssS variant strains showed marked heme sensitivity compared to native HssS-containing strain (Fig. 4C and Fig. S5A). We conclude that heme sensing is strongly dependent on Arg64 and Arg163, giving support to their role in anchoring heme as predicted by docking (Fig. 4).

**Figure 4.**
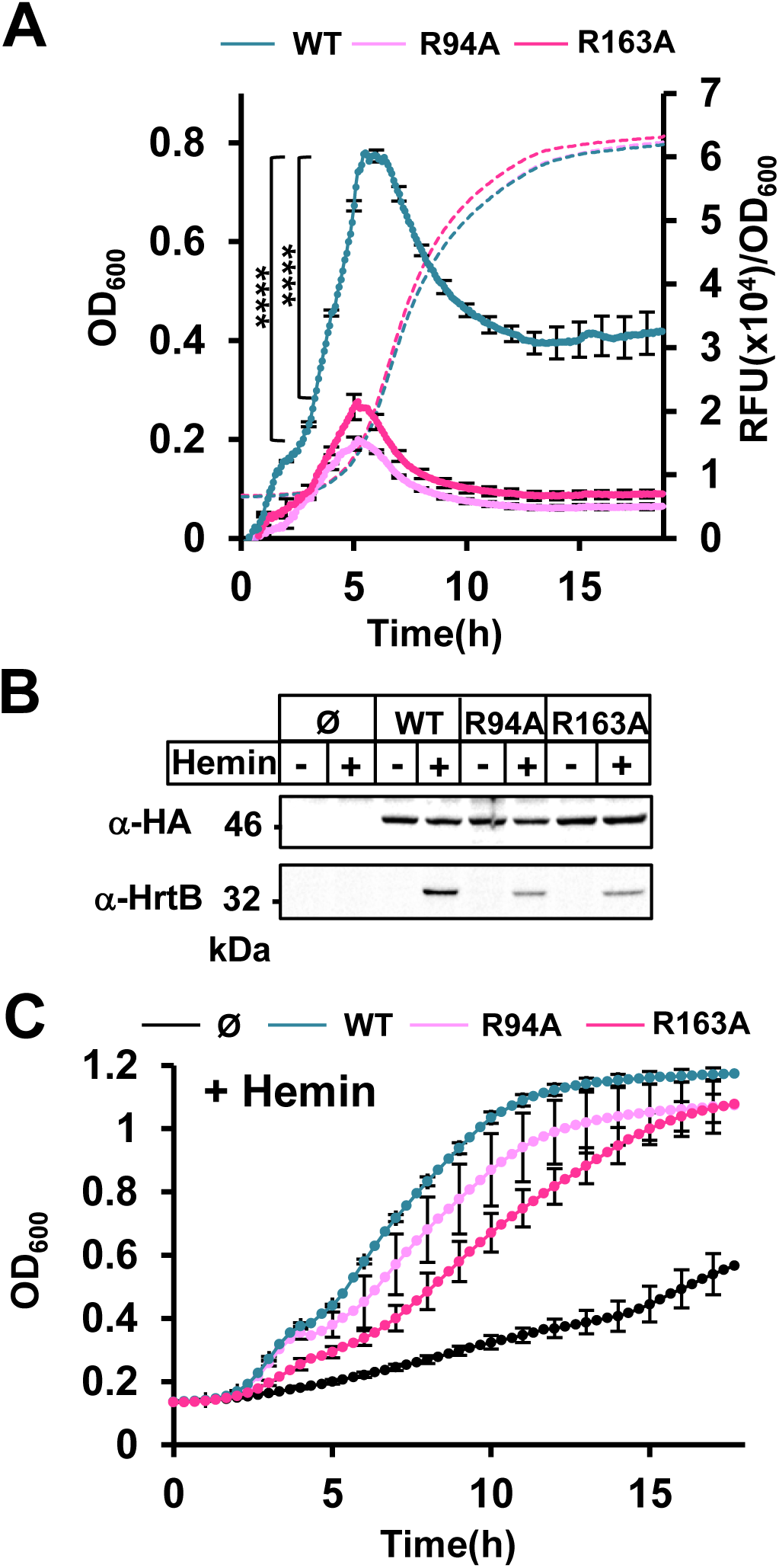
Pivotal roles of Arg94 and Arg163 on HssS activation. (A) P*_hrtBA_* transcriptional induction by HssS, HssS R94A and HssS R163A. Kinetics of P*_hrtBA_* induction in HG001 Δ*hssRS* mutant transformed either with pGFP(HssS), pGFP(HssS R94A) or pGFP(HssS R163A). Strains from ON cultures were diluted to OD_600_ = 0.01 in CDM ± 1 µM of hemin in a 96 well microplate. OD_600_ and GFP expression were followed in a spectrofluorimeter Infinite (Tecan) as in Fig. 2. Results of hemin induced fluorescence minus non-induced (background, 0 µM hemin) are displayed. Results represent the average ± S.D. from triplicate biological samples. ****, *P <* 0.0001, Student’s *t* test. The corresponding growth curves are shown. (B) Comparative expressions of HssS-HA, HssS-HA R94A, HssS-HA R163A and HrtB. Strains as in (A) were diluted in BHI from ON culture at OD_600_ = 0.01. Cultures were supplemented ± 1 µM hemin and grown for 1.5 h. HG001 Δ*hssRS* transformed with the empty plasmid (Ø) was used as a control. HssS-HA and HrtB expression were monitored on bacterial lysates by Western Blot (WB) with an anti-hemagglutinin antibody (α-HA) and an anti-HrtB antibody respectively (α-HrtB). Results are representative of three independent experiments. (C) Hemin toxicity in HssS, HssS R94A and HssS R163A expressing strains. Strains as in (B) were diluted from an ON preculture to an OD_600_ of 0.01 in BHI supplemented with 20 µM hemin and grown in a 96 microplate. (Growth curves were similar for all strains grown without hemin (Fig. S5A)). OD_600_ was recorded every 20 min for the indicated time in a spectrophotometer (Spark, Tecan). Results represent the average ± S.D from triplicate biological samples.

We next tested the importance of the predicted hydrophobic environment of heme. We choose phenylalanines, Phe25 and Phe128, which are predicted to be less than 4 Å from heme and could be engaged in π-π interactions that stabilize heme (Fig. 3E). The HG001 Δ*hssRS* mutant carrying either the F25A or F128A HssS variant (pGFP(HssS F25A) or pGFP(HssS F128A)) (Table S1) showed similar expression levels as the WT counterpart; however, both variants were defective for heme signal transmission to P*_hrtBA_* (Fig. 5A and B). Moreover, both variants showed increased heme sensitivity (Fig. 5C and Fig. S5B). As per predictions, Phe165, a conserved AA that is more distant from heme (Fig. 5D) and would only be able to contribute to heme stabilization from the edge of its aromatic ring, had a lower impact on HssS activation (Fig. 5D). We conclude that, analogous to Arg94 and Arg163A; Phe25, Phe128 are required for heme sensing and HssS function.

**Figure 5.**
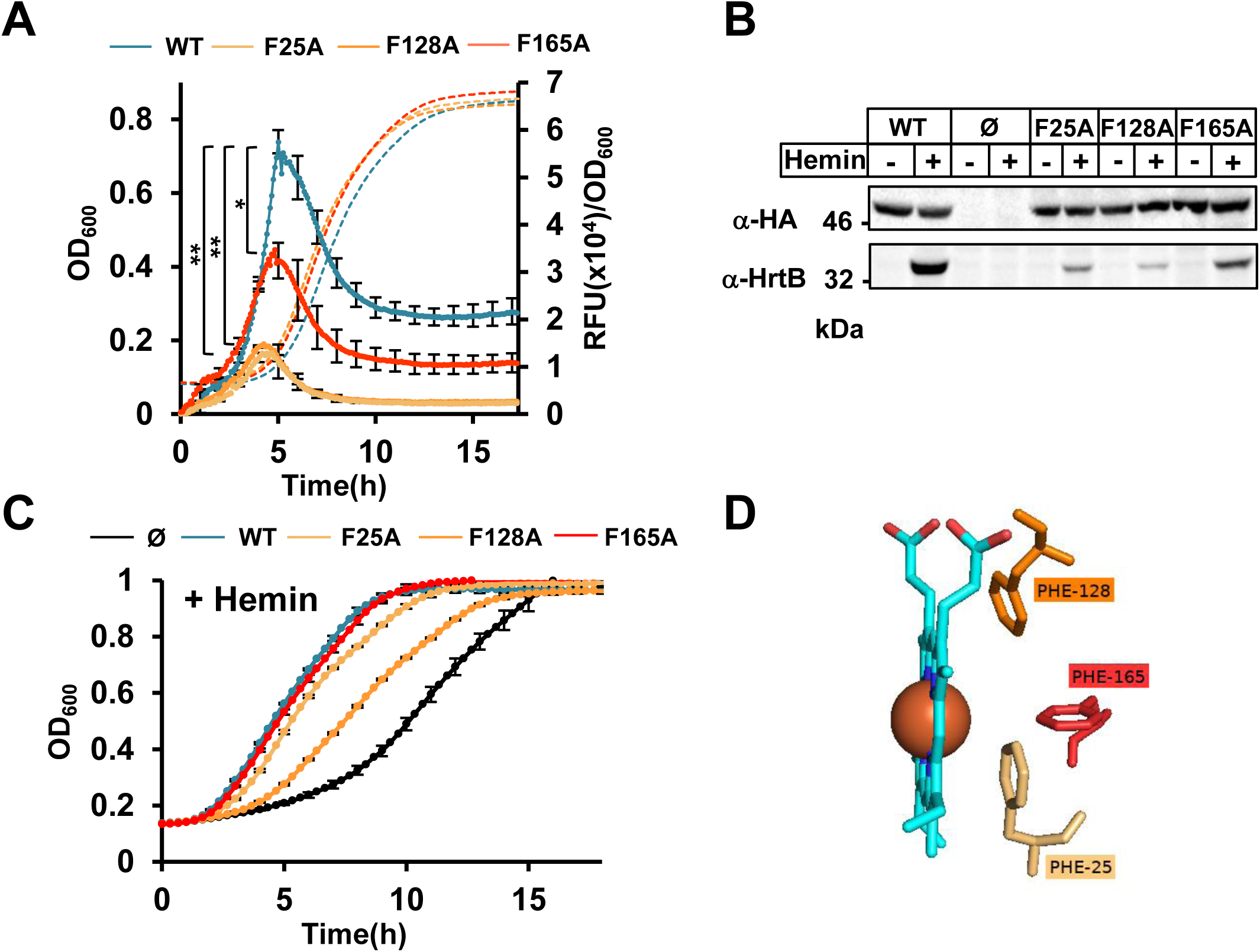
Impact of the heme binding hydrophobic environment on HssS activation. (A) P*_hrtBA_* transcriptional induction by HssS, HssS F25A, HssS F128A and HssS F165A. Kinetics of P*_hrtBA_* induction in HG001 Δ*hssRS* mutant transformed either with pGFP(HssS), pGFP(HssS R25A), pGFP(HssS F128A) or pGFP(HssS F165A) was performed as in Fig. 4. Results of hemin induced fluorescence minus non-induced (background, 0 µM hemin) are displayed. Results represent the average ± S.D. from triplicate biological samples. ****, *P <* 0.0001, Student’s *t* test. The corresponding growth curves are shown. (B) Comparative expressions of HssS-HA, HssS-HA F25A, HssS-HA F128A, HssS-HA F165A and HrtB. Strains as in (A) were diluted in BHI from ON culture at OD_600_ = 0.01. Cultures were supplemented ± 1 µM hemin and grown for 1.5 h. HG001 Δ*hssRS* transformed with the empty plasmid (Ø) was used as a control. HssS-HA and HrtB expression were monitored on bacterial lysates by Western Blot (WB) as in Fig. 4. Results are representative of three independent experiments. (C) Hemin toxicity in HssS, HssS F25A, HssS F128A and HssS F165A expressing strains. Strains as in (B) were diluted from an ON preculture to an OD_600_ of 0.01 in BHI supplemented with 20 µM hemin and grown in a 96 microplate as in Fig. 4. (Growth curves were similar for all strains grown without hemin (Fig. S5B). Results represent the average ± S.D from triplicate biological samples. (D) Pymol representation of the relative positions of F25, F128 and F165 to hemin in the predicted heme binding domain of HssS.

Finally, we constructed an HssS variant with the four mutations Arg64, Arg163, Phe25 and Phe128 (pGFP (HssS 4 mut.)) (Table S1), each of which is positioned at less than 4 Å from the porphyrin (Fig. 3E). Despite being expressed at WT levels, this variant was inactive and failed to induce HrtB expression (Fig. 6A and B). As expected, heme sensitivity of the strain expressing HssS 4 mut. was similar to that of the Δ*hssRS* strain in the presence of hemin (Fig. 6C and Fig. S5C).

**Figure 6.**
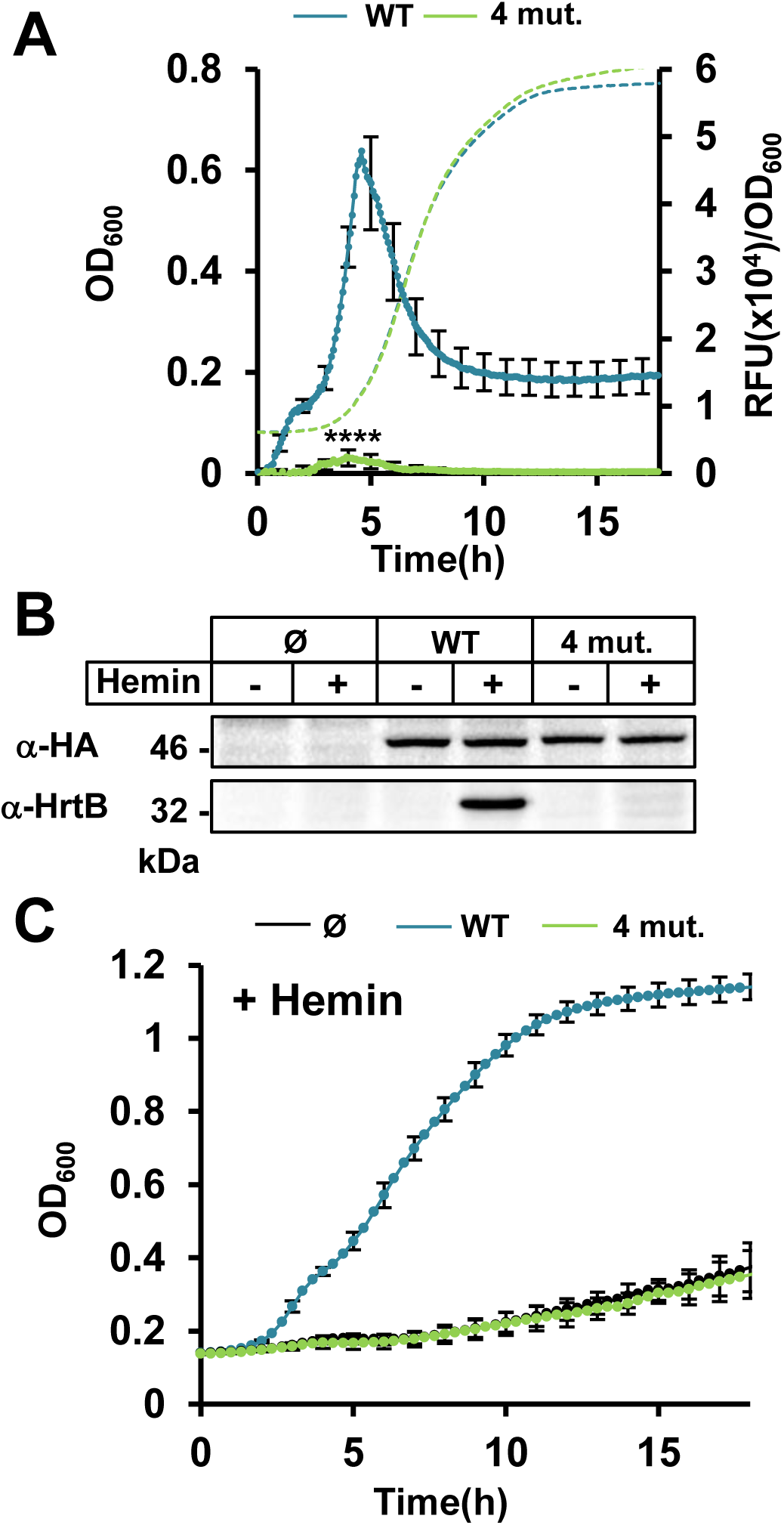
Inhibition of the heme sensing activity of HssS R94A, R163A, F25A, F128A (4 mut.). (A) HssS-4 mut. is unable to induce P*_hrtBA_* transcription. Kinetics of P*_hrtBA_* induction in HG001 Δ*hssRS* mutant transformed either with pGFP(HssS) or pGFP(HssS 4 mut.) was performed as in Fig. 4. Results of hemin induced fluorescence minus non-induced (background, 0 µM hemin) are displayed. Results represent the average ± S.D. from triplicate biological samples. ****, *P <* 0.0001, Student’s *t* test. The corresponding growth curves are shown. (B) Comparative expressions of HssS-HA, HssS-HA 4 mut. and HrtB. Strains as in (A) were diluted in BHI from ON culture at OD_600_ = 0.01. Cultures were supplemented ± 1 µM hemin and grown for 1.5 h. HG001 Δ*hssRS* transformed with the empty plasmid (Ø) was used as a control. HssS-HA and HrtB expression were monitored on bacterial lysates by Western Blot (WB) as in Fig. 4. Results are representative of three independent experiments. (C) Hemin toxicity in HssS and HssS 4 mut. expressing strains. Strains as in (B) were diluted from an ON preculture to an OD_600_ of 0.01 in BHI supplemented with 20 µM hemin and grown in a 96 microplate as in Fig. 4. (Growth curves were similar for all strains grown without hemin (Fig. S5C). Results represent the average ± S.D from triplicate biological samples.

Since the HssS variants tested are stable as shown by their expression on WB, we conclude that the predicted heme anchoring AAs and the integrity of the surrounding hydrophobic environment are essential for triggering HssS activation. These results supports the structural model of heme interaction with HssS and therefore question the role and importance of extracellular domain (ECD) of HssS.

### HssS lacking the extracellular domain [42-151] is constitutively activated

We examined the impact of removing most of the [35-168] domain of HssS corresponding to the ECD on heme signal transduction. A truncated version of *hssS* was constructed (referred to as pGFP(HssS ΔECD)) (Table S1), and was established in HG001 Δ*hssRS*. In this variant, the ECD AAs comprising AAs [35-41] and [151-168] were conserved and fused to allow membrane insertion and thus lacked Arg94 and Phe128 that are essential for heme sensing (see above). Expression and membrane localization of HssS-HA ΔECD were verified on WB using an anti-HA antibody following cell fractionation (Fig. S6A). Expression of the ECD variant compared to the full length protein was lower, possibly suggesting differences in protein stability (Fig. S6A).

To investigate the impact of the ECD deletion on HssS activity, GFP expression from pGFP(HssS) and pGFP(HssSΔECD) was followed in the absence or presence of 1 µM heme (Fig. 7 A and B). Remarkably, P*_hrtBA_*-GFP was expressed constitutively and independently of heme addition in the strain carrying pHssS ΔECD (Fig. 7 A and B). However, while HrtB expression was constitutive, its levels were lower than in the strain producing WT HssS (Fig. 7C and Fig. S6A). Interestingly, despite lower levels of HrtB in the strain expressing HssSΔECD, the strain showed markedly improved fitness when challenged with 10 µM heme when compared to the isogenic strain expression HssS WT (Fig. S6B and C). This observation appears to indicate a fitness advantage during infection of bacteria carrying a defective HssS protein, such that *hrtBA* is constitutively active.

**Figure 7.**
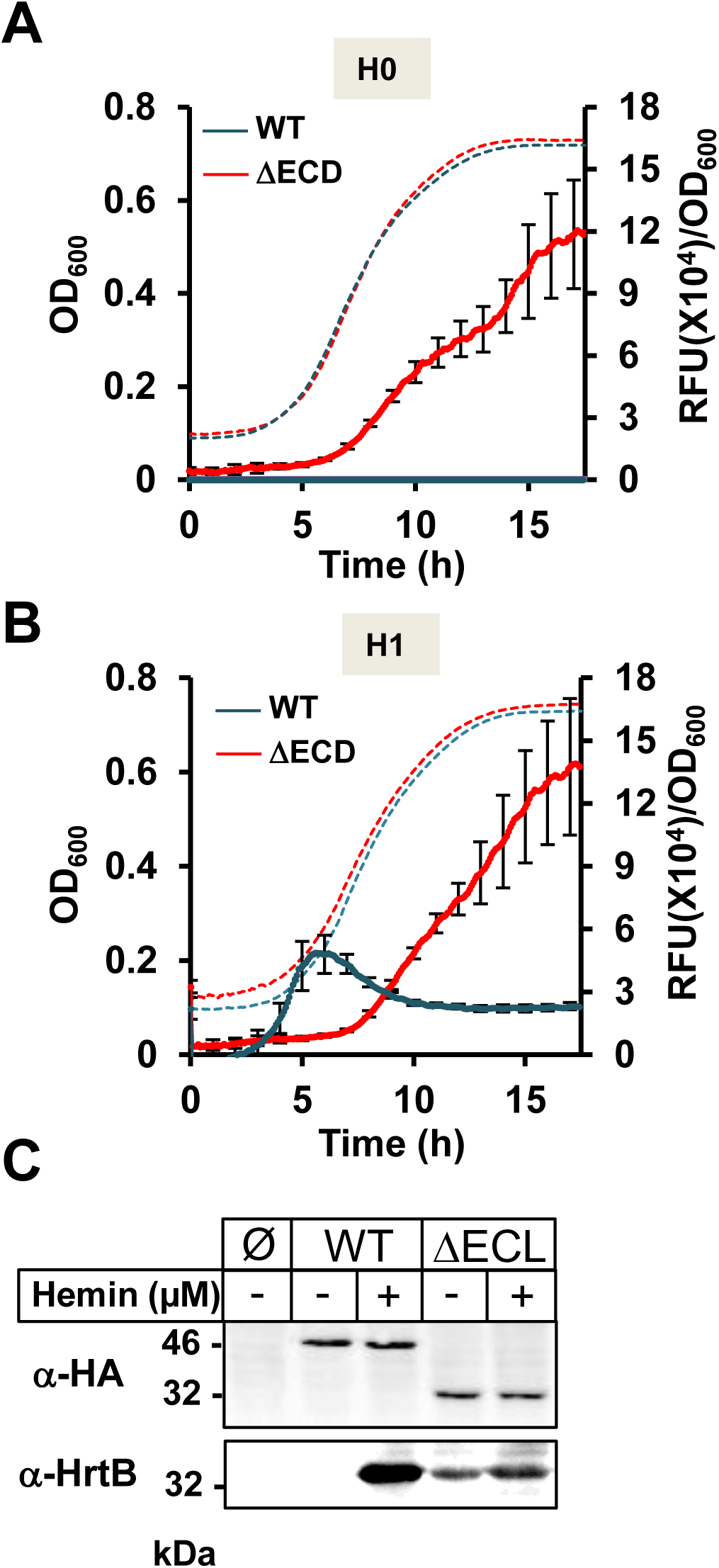
The extracellular domain is required for heme dependent activation of HssS. (A-B) Comparative dynamics of P*_hrtBA_* transcriptional induction by HssS WT and HssS ΔECD HG001 in absence (A) or in presence of hemin (B). HG001 Δ*hssRS* mutant transformed with either pGFP(HssS), pGFP(HssS ΔECD) or empty vector were grown as in Fig. 2 in CDM without heme (A) or supplemented with 1 µM of hemin (B). Fluorescence (RFU) and OD_600_ were determined as in Fig. 2. Results of fluorescence (RFU/OD_600_) from strains expressing HssS WT or HssS ΔECD minus empty vector transformed strain are displayed. Results represent the average ± S.D. from triplicate biological samples. The corresponding growth curves are shown. (C) HssS ΔECD constitutively signals HrtB expression. Δ*hssRS* HG001 transformed either with pGFP(HssS), pGFP(HssS ΔECD) or empty vector (Ø) were used to monitor HssS and HrtB expression by WB with α-HA and an α-HrtB, respectively. Bacteria were grown to an OD_600_=0.5 and induced for 1.5 h ± 1 µM hemin. SDS-PAGE was performed on cell lysates (25 μg per lane). Results are representative of three independent experiments.

We conclude that in the absence of ECD, HssS does not retain its capacity to be stimulated by heme, but also appears to transmit a basal signal leading to *hrtBA* induction and bacterial protection against heme. The HssS ΔECD phenotype contrasts with that of the Δ*hssS* strain, which fails to induce *hrtBA*. It is thus tempting to speculate that HssS ΔECD maintains the heme binding domain open to accommodate heme. Altogether, these data point out the importance of the extracellular domain for HssS activation.

## Discussion

Heme homeostasis in Gram positive bacteria is mainly achieved *via* heme efflux. In *S. aureus*, the major heme efflux transporter HrtBA is controlled by HssRS (13, 18, 27). Our results give strong support that the heme sensing function of the dimeric HK HssS is located in a structural domain at the interface between the TM and extracellular domains. Thus, membrane-attached rather than extracellular heme pools control HssS transient activation, shedding new light on a detoxification mechanism of an abundant host molecule in a major human pathogen *S. aureus*.

Four lines of evidence support the hydrophobic HssS interface as being required for heme binding: 1-In depth *in silico* modelling predicted HssS protein structure and heme binding candidate amino acids. In heme docking simulations, heme interacted with both periplasmic and lipid-embedded amino acids. 2-Loss of functional HssS activity correlated with directed mutagenesis of heme binding candidate amino acids, consistent with their functional roles. Mutations of targeted AA in the hydrophobic environment and in predicted anchoring arginines (Arg94 and Arg163), were all required to abolish HssS activation. 3-The implicated amino acids and overall structure of *S. aureus* HssS are conserved in protein homologs in other bacteria. These findings also clarify previous work in which random mutations of conserved AA residues required for HssS heme sensing in *S. aureus* and in *Bacillus anthracis* were mapped to the same domain (19, 20). 4-Heme availability from the outside or from the inside lead to similar kinetics of induction. HssS is activated by extracellular heme, but also by increasing intracellular heme pools (*e.g.*, in a Δ*hrtBA* mutant), suggesting that both heme sources are accessible to HssS binding; by deduction, this common site would need to be the membrane.

Our findings implicate heme bound to the HssS membrane-outer surface interface as activation signal of HssS. We suggest that this mechanism of TCS activation may more generally be a novel basis for hydrophobic molecule sensing. A previously reported class of HKs called intramembrane histidine kinase (IM-HK) perceives its stimuli in the membrane, but not directly (28–30). Instead, an N-terminal signal transfer region consisting of two transmembrane helices presumably connects the IM-HKs with the regulated accessory membrane proteins that function as the true sensors. For HKs that lack most of the sensory domain in the ECD, cell envelope stress sensing has been directly linked to the ABC transporter *via* TM-TM interactions (28–30). In contrast, an activation mechanism based on direct physical interaction between HssS and HrtBA seems unlikely since P*_hrtBA_* is induced in an *hrtBA* deletion mutant. HssS is thus not activated similarly to IM-HK. We thus propose HssS as a paradigm for signaling by organic molecules, as produced by the host, which are in contact with the membrane-surface interface. Membrane input could be particularly relevant for regulators of efflux transporters controlling exogenous substrates, including antibiotics (31–33).

*In silico* simulation revealed the membrane-surface interface as the site of HssS interaction with heme, but does not take into account the role of the flexible phospholipid environment, which varies according to conditions and environmental lipids (34). Since intramembrane heme concentrations impact HssS activation (as seen by testing HssS induction in the Δ*hrtBA* mutant), it is highly likely that heme crosses the membrane to activate HssS. Membrane lipid properties would then be expected to modulate HssS activation. We are currently investigating the impact of altering phospholipid composition on HssS expression to test this prediction.

Our findings that membrane heme activates HssS provides a functional link between HssS and HrtBA. HrtB is a member of the MacB family of efflux pumps that is distinct from other structurally characterized ABC transporters (35). The structural basis for heme efflux by HrtB was recently solved in *Corynebacterium diphteriae*: the HrtB dimer forms a heme binding site in the outer leaflet of the membrane, which is laterally accessible to heme (17). HssS would have the integral role as heme “gatekeeper” that controls exogenous heme pools to prevent translocation within the membrane and into the cytosol (Fig. 8). Membrane heme may either enter passively into the intracellular compartment or be effluxed by HrtB before this step. This alternative model is compatible with our observations that exogenous excess heme is internalized in *S. aureus* independently of the Isd heme import system in our experimental conditions ((36) and data not shown).

**Figure 8.**
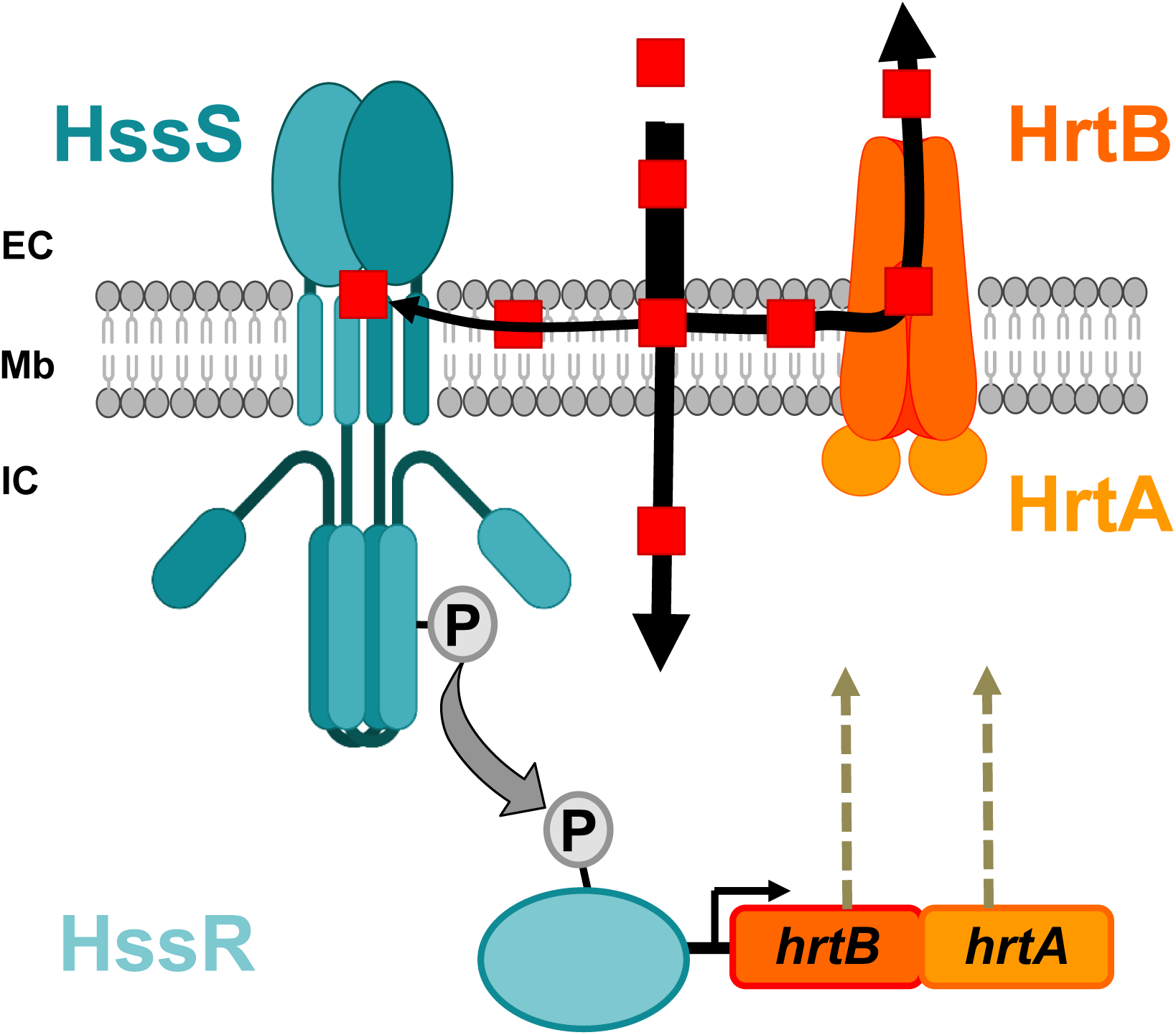
Functional model of exogenous heme management by the gatekeeping HssRS-HrtBA system in *S. aureus.* Exogenous hemin (red dots) translocates through the membrane (Mb) compartment from the extracellular (EC) to the intracellular (IC) compartments. HssS senses heme at the membrane interface, activating the phosphorelay between the HK and HssR leading to the expression of HrtBA. The pool of hemin that crosses the membrane by diffusion is counterbalanced by HrtBA which extrudes heme from the membrane to EC space (17).

Unlike *S. aureus*, *L. lactis* and *E. faecalis* control HrtBA expression using intracytoplasmic TetR transcriptional regulators (HrtR in *L. lactis* (12) and FhtR in *E. faecalis* (10)). It is thus tempting to speculate that possible that HssS sensing mechanism discriminates heme originating from endogenous synthesis from exogenous sources, which would minimize interference with metabolic processes.

*Staphylococcus aureus* is a serious threat to public health due to the rise of antibiotic resistance in this organism. As such, many efforts are under way to develop therapies that target essential host adaptive processes in *S. aureus*. Our findings provide a new basis for the elucidation of pathogen sensing mechanisms as a prerequisite to the discovery of inhibitors.

## Materials and methods

### Bacterial strains

The strains and plasmids used in this work are listed in supplemental Table S1. Plasmid constructions procedures are outlined in the supplemental data. *Staphylococcus aureus* strain HG001, a derivative of the RN1 (NCT8325) strain with restored *rbsU* (a positive activator of SigB) (37). HG001 Δ*hrtBA* and Δ*hrssS* mutants construction are described in supplemental information section.

### Bacterial Growth Conditions and Media

*Staphylococcus aureus* HG001 and USA300 and their derivatives were grown as ON pre-cultures at 37°C in rich BHI liquid broth (DIFCO) supplemented with 0.2 % glucose with aeration by shaking at 200 rpm. All growth assays were performed in a 96-well plate in 200 µl of BHI. Optical density at 600 nm (OD_600_) served as a measurement of growth and was measured every 15 min for the indicated total time in a microplate reader (Spark, Tecan). *E. coli* strains were grown in Luria-Bertani (LB) medium at 37°C with aeration by shaking at 180 rpm. When needed, antibiotics were used as follows: 50 µg/ml kanamycin and 100 µg/ml ampicillin for *E. coli*; 5 µg/ml erythromycin for *S. aureus*. Hemin was prepared from a stock solution of 10 mM hemin chloride dissolved in 50 mM NaOH; Frontier Scientific).

### Dynamics of fluorescence emission

For kinetic studies using GFP, *S. aureus* strains were grown a 96-well plate in 200 µl CDM (chemically defined medium) as reported (38, 39). CDM medium contained around 170 nM iron (38). CDM is uncolored therefore minimizing fluorescence background. OD_600_ and fluorescence (Exc.: 480 nm; Em.: 515 nm, bandwidth: 9 nm, integration time: 40 µs, gain: 140) were measured every 5 min in a black 96-well microplate with transparent bottom (Greiner Bio-one, Kremsmünster, Austria) in a spectrofluorimeter (Infinite M200, Tecan) at 37 °C under constant shaking (orbital, amplitude: 2.5).

### Antibodies

An anti-HrtB antibody targeted to the extracellular domain of *S. aureus* HG001 HrtB was produced. The HrtB[45-236] fragment was purified from *E. coli* as a His-tagged antigen from the plasmid pET200-*hrtB* ECL (see above and Table S1), purified from bacterial lysates on a nickel affinity resin (His-Select, Sigma-Aldrich) as described (12). Briefly, *E. coli* BL21 (DE3) (Thermo Fisher France, Villebon-sur-Yvette) transformed with pET200-*hrtB* ECL was grown to OD_600_ = 0.6, and expression was induced with 1 mM isopropyl 1-thio-β-d-galactopyranoside for 2 h at 37 °C. Cells were pelleted at 3,500 × *g* for 10 min, resuspended in 50 mm Tris-HCl, pH 8.0, 300 mm NaCl, containing 20 mm imidazole (binding buffer), and disrupted with glass beads (Fastprep, MP Biomedicals France, Illkirch-Graffenstaden). Cell debris were removed by centrifugation at 18,000 × *g* for 15 min at 4 °C. The soluble fraction (surpernatant) was mixed with the nickel affinity resin and incubated on a spinning wheel at 4°C for 1 h. The resin was then centrifuged (5000 rpm, 5 min) and washed three times with binding buffer. Purified proteins were eluted with 50 mm Tris-HCl, pH 8.0, 300 mm NaCl, containing 150 mm imidazole, dialyzed against 50 mm Tris-HCl, pH 7.5, and finally stored at −80 °C. Protein concentrations were determined with the Lowry assay method (Bio-Rad France, Marnes-la-Coquette). The resulting purified His-HrtB-ECL was used for rabbit antibody production (Covalab, Bron, France). Antiserum specificity was determined by Western blots using known amounts of purified His-HrtB-ECL proteins and bacterial lysates expressing HrtB. The polyclonal anti-GAPDH antibody was a generous gift from F. Götz (University of Tübingen, Germany) (40). The polyclonal anti-HA antibody was from Thermo Fisher.

### Bacterial lysate and membrane isolation

Bacteria were pelleted by centrifugation and washed with PBS. Resuspended cells were then pelleted at 3,500 g for 10 min, resuspended in 50 mM Tris-HCl pH 7.5, 150 mM NaCl, containing 0.2% Triton (lysis buffer), and disrupted with glass beads (Fastprep; MP Biomedicals). Cell debris were removed by centrifugation at 18,000 *g* for 15 min at 4°C. To prepare membranes, bacterial lysates were prepared as above except that bacteria were lysed in 20 mM Tris-HCl pH 7.5. Lysates were then submitted to centrifugation at 100,000 *g* for 45 min at 4°C in an Ultracentrifuge Beckman XL-90 (Beckman France, Villepinte) equipped with a 70.1T1 rotor. Membranes pellets were resuspended in lysis buffer. Proteins were quantified by the Lowry Method (Bio-Rad) and denatured in Laemmli sample buffer at 95°C for 5 min.

### β-galactosidase assays

β-galactosidase activity was quantified by luminescence in an Infinite M200 spectrolumineter (Tecan) using the luminescence β-glo assay as recommended by the manufacturer (Promega France, Charbonnières-les-Bains, France). Briefly, *S. aureus* strains cultures were diluted from ON precultures in BHI to an OD_600_ = 0.01 and grown to an OD_600_ = 0.5 and then incubated for 1 h with the indicated concentrations of hemin. 25 µl cultures were distributed in a white 96-wells microplate (Greiner Bio-one) in triplicate. 50 µl β-glo assay reagent was added per well. After 10 min incubation at RT, luminescence was quantified. In parallel, 200 µl of the corresponding cultures were distributed in a transparent 96-wells plate to measure the OD_600_ for normalization of the luminescence.

### Heme concentration determination in bacterial lysates

Proteins from bacterial lysates (as described above) (in a volume of 250 µl) were mixed with 20 µl of 0.2 M NaOH, 40 % (v/v) pyridine and 500 µl potassium ferricyanide or 5 mM sodium dithionite. 500-600 nm absorption spectra were recorded in a UV-visible spectrophotometer Libra S22 (Biochrom, Cambridge,UK). Dithionite-reduced minus ferricyanide-oxidized spectra of pyridine hemochromes were used to determine the amount of heme *b* by following the value of the difference between absorbance at 557 nm (reduced) and 540 nm (oxidized) using a difference extinction coefficient of 23.98 nM^-1^.cm^-1^ (41).

### Heme docking analysis

Heme (https://www.rcsb.org/) was docked onto the modeled HssS structure with AutoDock Vina 1.1.2 (https://ccsb.scripps.edu/). The exhaustivity parameter was set to 8. Ligand and protein coordinates were prepared (including polar hydrogen atoms and atoms charges to take hydrogen and electrostatic interactions into account) using Open babel (http://openbabel.org/wiki/Main_Page). Docking of heme was performed on the entire surface of one monomer in the dimeric HssS with 100 boxes (27.10^-33^.m^-3^). From each box, the top scoring pose (in terms of binding free energy (kcal/mol) as estimated by AutoDock Vina) was selected for binding site. The 10 top docking solutions were visualized with VMD ((42), http://www.ks.uiuc.edu/Research/vmd/) and designated a single periplasmic.

## Abbreviations

AF: AlphaFold
β-gal: β-galactosidase
CDM: Chemically Defined Medium
ECD: Extracellular Domain
HAMP: Histidine kinases, Adenyl cyclases, Methyl-accepting proteins and Phosphatases
Hb: hemoglobin
HK: Histidine Kinase
HisKA: family A of histidine kinase
Hrt: Heme-regulated transport
HssS: Heme sensing system Sensor
HssR: Heme sensing system Regulator
HrtR: Heme-regulated transport Regulator
IM-HK: Intramembrane Histidine Kinase
OPM: Orientation of Proteins in Membranes
PDC: PhoQ-DcuS-CitA
RFU: Relative Fluorescence Unit
RLU: Relative Luminescence Unit
TCS: Two-Component System
TM: Transmembrane
WB: Western blot

## Acknowledgments

This work was supported by the HemeDetox - 17-CE11-0044-01 project by the French “Agence Nationale de la Recherche”. V. Saillant is the recipient of a doctoral fellowship from the French ministry of Research and Paris-Saclay University. The funders had no role in study design, data collection and analysis, decision to publish, or preparation of the manuscript. The funders had no role in study design, data collection and analysis, decision to publish, or preparation of the manuscript. We thank Dr. A. Gruss (INRAE, France) and Dr. P. Delepelaire (IBPC, France) for their technical help and insightful comments on our work. We are grateful to A. Hiron (Université de Tours, France) and T. Msadek (Institut Pasteur, France) for the HG001 Δ*hssRS Staphylococcus aureus* strain and to Dr. F. Götz (University of Tübingen, Germany) for the generous gift of the anti-GAPDH antibody.

## Supplemental Information

### Plasmid construction

Plasmid pTCV-*lac* is a low copy number plasmid that uses the *lac* gene as a reporter to evaluate promoter activities in Gram-positive bacteria (43) (*SI Appendix*, Table S1). DNA fragments containing the *hrtBA* or the *hssRS* promoter were PCR-amplified with primer pairs (O1-O2) or (O3-O4) (*SI Appendix*, Table S2) respectively. The amplified fragments were digested with *Eco*RI/*Bam*HI and cloned into pTCV-*lac*, resulting in plasmids pP_hrtBA_-*lac* and pP_hssRS_-*lac* (*SI Appendix*, Table S1). pP*_hrtBA_*-GFP (*SI Appendix*, Table S2) was constructed by cloning fragments corresponding to P*_hrtBA_* with the primer pairs (O5-O6) and into the *Bam*H1/*Eco*R1 restriction sites of the pCN52 vector (*SI Appendix*, Table S1) using HG001 genomic DNA as template. The DNA sequence of P_hssRS_ *hssRS*-HA was PCR amplified from the plasmid pUC *hssRS*-HA-P_hrtBA_ with oligonucleotides (O7-O8) (*SI Appendix*, Table S2), digested with *Bam*H1/*Kpn*1 and ligated into pCN52 to give rise to p*hssRS*-HA, P*_hrtBA_*-GFP (pGFP(HssS)) (*SI Appendix*, Table S1). *hssS* gene is followed by the nt sequence encoding the hemagglutinin influenza epitope (HA, YPYDVPDYA). p*hssRS*-HA ΔECL, P*_hrtBA_*-GFP (pGFP (HssS ΔECL)) was obtained by an overlap of two PCRs amplified with the primer pairs (O7-O9) and (O8-O10) (*SI Appendix*, Table S2) using pUC *hssRS*-HA-P*_hrtBA_* as a template. The P*_hssRS_hssRS*-HA ΔECL fragment digested with *Bam*H1/*Kpn*1 and ligated into pCN52 to give rise to pGFP (HssS ΔECL) (*SI Appendix*, Table S1). The DNA sequence corresponding to extracellular domain of HrtB was PCR amplified with primers pair (O11-O12) using HG001 genomic DNA as a template that was cloned into the *E. coli* expression vector pET200 as recommended by the manufacturer’s instructions to generate pET200-*hrtB* ECL (*SI Appendix*, Table S1). The pMADΔ*hrtBA* plasmid was constructed as follows: 2 DNA fragments of ∼800 bp flanking the *hrtBA* operon were PCR-amplified using *S. aureus* HG001 genomic DNA as a template and primer pairs (O13-O14) and (O15-O16) (*SI Appendix*, Table S2). Both fragments were used as templates in a second round of PCR amplification with primers (O13-O16), resulting in an overlapping ∼1.6 kb fragment, which was digested by *Xma*I and *Bam*HI and cloned into the thermosensitive plasmid, pMAD (2), giving rise to pΔ*hrtBA* (*SI Appendix*, Table S1). pΔ*hrtBA* was established by transformation in *S. aureus* HG001. The double cross-over event leading to the Δ*hrtBA* mutant was obtained as described (44). p*hssRS*-HA was constructed by PCR amplification of the hssRS-HA sequence from the pUC *hssRS*-HA-P*_hrtBA_* plasmid with (O16-O17). The amplified P*_hssRS_ hssRS*-HA DNA fragment was cloned into *Pst*1/*Bam*H1 restriction sites of PAW8 (*SI Appendix*, Table S1). All plasmids were verified by DNA sequencing.

1. Poyart C, Trieu-Cuot P. 1997. A broad-host-range mobilizable shuttle vector for the construction of transcriptional fusions to B-galactosidase in Gram-positive bacteria. FEMS Microbiology Letters 156:193-198.
2. Arnaud M, Chastanet A, Debarbouille M. 2004. New vector for efficient allelic replacement in naturally nontransformable, low-GC-content, gram-positive bacteria. Appl Environ Microbiol 70:6887-91.

### Supplemental Tables

**Table S1.** Strains and plasmids used in this study.

| *Strain/plasmid* | *Characteristics* | *Source/*<br>*reference* |
| --- | --- | --- |
| **Strain** |  |  |
| ***E. coli*** |  |  |
| TOP10 | *F<sup>-</sup> mcrA Δ(mrr-hsdRMS-mcrBC) Δ80lacZΔM15 lacX74 recA1 deoR araD139 Δ(ara-leu)7697 galU galK rpsL (Str<sup>r</sup>) endA1* | Invitrogen |
| BL21 (DE3) | *lacI<sup>q</sup> rrnB T14 ΔlacZΔWJ16 hsdR514 ΔaraBA-D<sub>AH33</sub> ΔrhaBAD<sub>LD78</sub>* | Invitrogen |
| ***S. aureus*** |  |  |
| RN4220 | *S. aureus* cloning recipient ATCC 8325-4 derivative restriction negative | (45) |
| HG001 | *Staphylococcus aureus* HG001 strain, derivative of the RN1 (NCT8325) strain with restored rbsU (a positive activator of SigB). | (37) |
| HG001H1 | HG001 Δ*hssRS*, deletion of *hssR* and *hssS* genes | This study |
| HG001H2 | HG001 Δ*hrtBA*, deletion of *hrtB* and *hrtA* genes | This study |
| **Plasmid** |  |  |
| pTCV-*lac* | Conjugative *E. coli* Gram-positive bacteria shuttle plasmid carrying the promoter less *E. coli lacZ* gene for constructing transcriptional fusions. Kan<sup>R</sup>, Ery<sup>R</sup> | (43) |
| pCN52 | *E. coli-S.aureus* bacteria shuttle plasmid with *gfpmut2* reporter construct. Amp<sup>R</sup>, Ery<sup>R</sup> | (46) |
| pAW8 | Cloning shuttle vector, pMB1 ori for replication in *E. coli*, pAMα1 ori for replication in gram-positive organisms. Tet<sup>R</sup>, Amp<sup>R</sup> | (47) |
| pMAD | Cloning shuttle replication-thermosensitive vector for generating stable chromosomal mutations in low G-C Gram positive bacteria. Amp<sup>R</sup>, Ery<sup>R</sup> | (44) |
| pP<sub>hrtBA</sub>-*lac* | *S. aureus* HG001 *hrtBA* promoter region cloned into pTCV-*lac*. Kan<sup>R</sup>, Ery<sup>R</sup> | This study |
| pP<sub>hssRS-lac</sub> | *S. aureus* HG001 *hssRS* promoter region region cloned into pTCV-*lac*. Kan<sup>R</sup>, Ery<sup>R</sup> | This study |
| pUC *hssRS*-HA, P<sub>hrtBA</sub> | P<sub>hssRS</sub> promoter region and *hssRS* genes followed by the promoter region of *hrtBA* from *S. aureus* HG001 cloned into pUC. Amp<sup>R</sup> | Proteogenix (France) |
| p*hssRS*-HA | *hssRS* promoter region and *hssRS* genes with a HA epitope at the Ct of *hssS* from *S. aureus* HG001 in pAW8. Tet<sup>R</sup>, Amp<sup>R</sup> | This study |
| pP<sub>hrtBA-gfp</sub> | *S. aureus* HG001 *hrtBA* promoter region cloned into pCN52. Amp<sup>R</sup>, Ery<sup>R</sup> | This study |
| pGFP(HssS) | p*hssRS*-HA, P<sub>hrtBA-gfp</sub>. Promoter region and *hssRS* genes with a HA epitope at the Ct of *hssS* from *S. aureus* HG001 and promoter region of *hrtBA* cloned upstream GFP in pCN52. Amp<sup>R</sup>, Ery<sup>R</sup> | This study |
| pGFP(HssS T253A) | p*hssRS*-HA T253A, P<sub>hrtBA-gfp</sub>. pGFP(HssS) plasmid with a mutation of the codon encoding T253 to alanine codon in *hssS*. Amp<sup>R</sup>, Ery<sup>R</sup> | E-Zyvec (France) |
| pGFP(HssS R94A) | p*hssRS*-HA R94A, P<sub>hrtBA-gfp</sub>. pGFP(HssS) plasmid with a mutation of the codon encoding R94 to alanine codon in *hssS*. Amp<sup>R</sup>, Ery<sup>R</sup> | E-Zyvec (France) |
| pGFP(HssS R163A) | p*hssRS*-HA R163A, P<sub>hrtBA-gfp</sub>. pGFP(HssS) plasmid with a mutation of the codon encoding R163 to alanine codon in *hssS*. Amp<sup>R</sup>, Ery<sup>R</sup> | E-Zyvec (France) |
| pGFP(HssS R94A, R163A) | p*hssRS*-HA R94A, R163A, P<sub>hrtBA-gfp</sub>. pGFP(HssS) plasmid with mutations of the codon encoding R94 and R163 to alanine codon in *hssS*. Amp<sup>R</sup>, Ery<sup>R</sup> | E-Zyvec (France) |
| pGFP(HssS F25A) | p*hssRS*-HA F25A, P<sub>hrtBA-gfp</sub>. pGFP(HssS) plasmid with a mutation of the codon encoding F25 to alanine codon in *hssS*. Amp<sup>R</sup>, Ery<sup>R</sup> | E-Zyvec (France) |
| pGFP(HssS F128A) | p*hssRS*-HA F128A, P<sub>hrtBA-gfp</sub>. pGFP(HssS) plasmid with a mutation of the codon encoding F128 to alanine codon in *hssS*. Amp<sup>R</sup>, Ery<sup>R</sup> | E-Zyvec (France) |
| pGFP(HssS F165A) | p*hssRS*-HA F165A, P<sub>hrtBA-gfp</sub>. pGFP(HssS) plasmid with a mutation of the codon encoding F165 to alanine codon in *hssS*. Amp<sup>R</sup>, Ery<sup>R</sup> | E-Zyvec (France) |
| pGFP(HssS4 mut.) | p*hssRS*-HA R94A, R163A, F128A, F165A, P<sub>hrtBA-gfp</sub>. pGFP(HssS) plasmid with mutations of the codon encoding R94, R163, F25, F128 to alanine codon in *hssS*. Amp<sup>R</sup>, Ery<sup>R</sup> | E-Zyvec (France) |
| pGFP(HssS ΔECL) | p*hssRS* ΔECL-HA, P<sub>hrtBA-gfp</sub>. pGFP(HssS) plasmid with a deletion of the nucleotides corresponding to the AA [50-129] of HssS. Amp<sup>R</sup>, Ery<sup>R</sup> | This study |
| pΔ*hssRS* | *hssRS* fragment cloned into pMAD to obtain the HG001-Δ*hssRS* mutant. | (48) |
| pΔ*hrtBA* | *hrtBA* fragment cloned into pMAD to obtain the HG001 Δ*hrtBA* mutant. Amp<sup>R</sup>, Ery<sup>R</sup> | This study |

1. Kreiswirth BN, Lofdahl S, Betley MJ, O’Reilly M, Schlievert PM, Bergdoll MS, Novick RP. 1983. The toxic shock syndrome exotoxin structural gene is not detectably transmitted by a prophage. Nature 305:709-12.
2. Caldelari I, Chane-Woon-Ming B, Noirot C, Moreau K, Romby P, Gaspin C, Marzi S. 2017. Complete Genome Sequence and Annotation of the Staphylococcus aureus Strain HG001. Genome Announc 5.
3. Poyart C, Trieu-Cuot P. 1997. A broad-host-range mobilizable shuttle vector for the construction of transcriptional fusions to B-galactosidase in Gram-positive bacteria. FEMS Microbiology Letters 156:193-198.
4. Charpentier E, Anton AI, Barry P, Alfonso B, Fang Y, Novick RP. 2004. Novel cassette-based shuttle vector system for gram-positive bacteria. Appl Environ Microbiol 70:6076-85.
5. Wada A, Watanabe H. 1998. Penicillin-binding protein 1 of Staphylococcus aureus is essential for growth. J Bacteriol 180:2759-65.
6. Arnaud M, Chastanet A, Debarbouille M. 2004. New vector for efficient allelic replacement in naturally nontransformable, low-GC-content, gram-positive bacteria. Appl Environ Microbiol 70:6887-91.
7. Toledo-Arana A, Merino N, Vergara-Irigaray M, Debarbouille M, Penades JR, Lasa I. 2005. *Staphylococcus aureus* develops an alternative, ica-independent biofilm in the absence of the arlRS two-component system. J Bacteriol 187:5318-29.

**Table S2.**
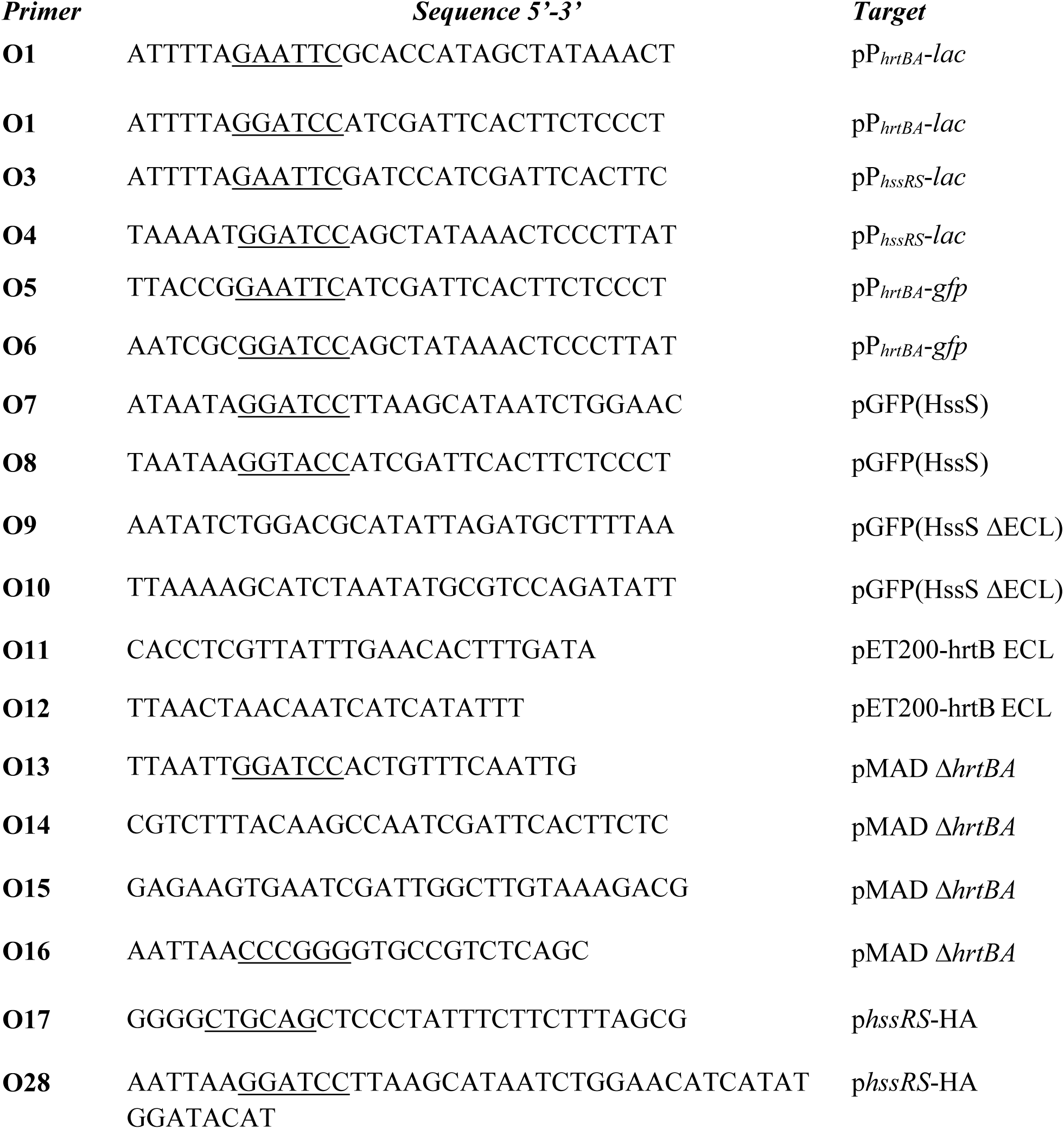
Oligonucleotides used in this study.

### Supplemental figures

**Figure S1.**
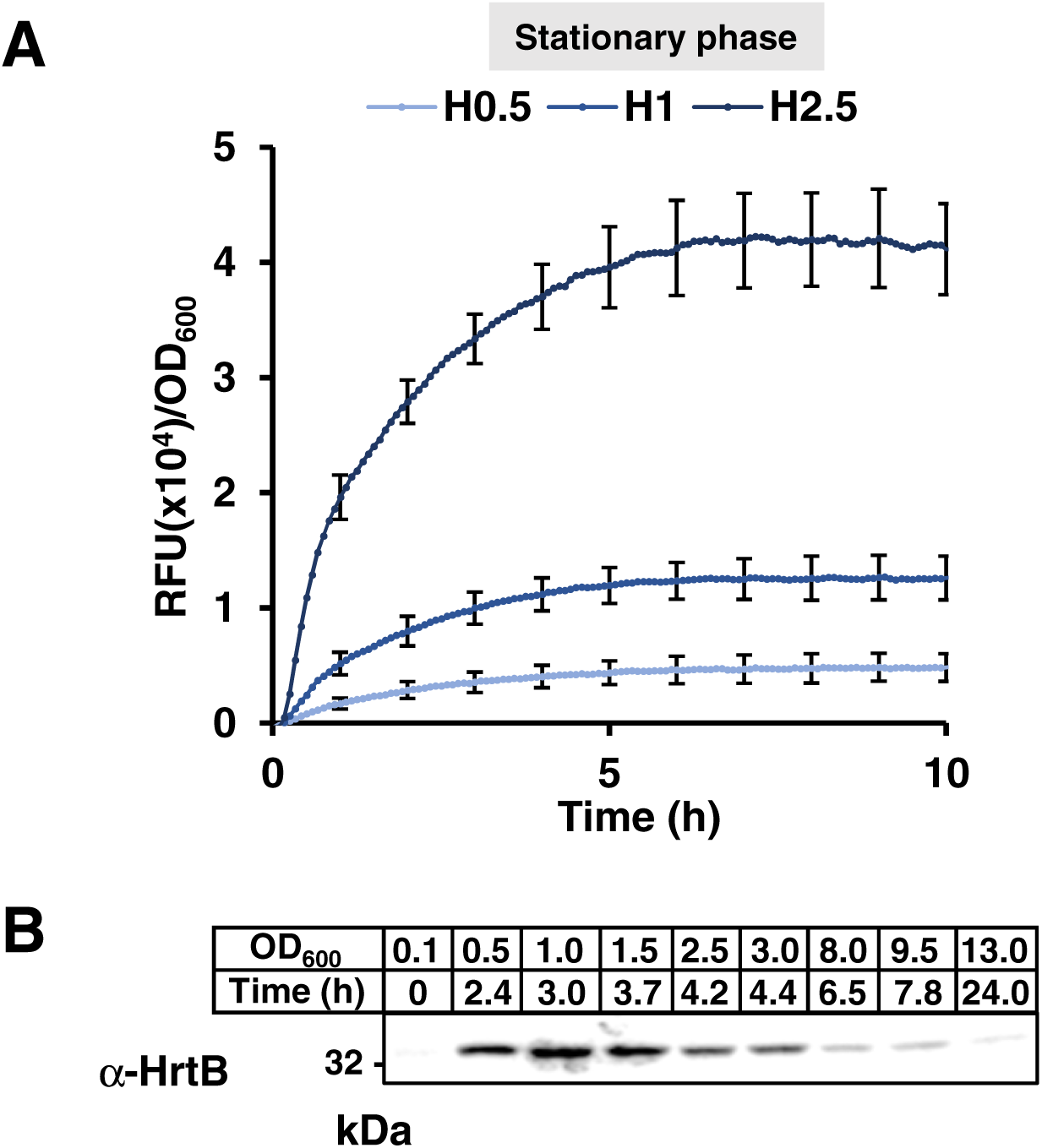
Transient induction of P*_hrtBA_* during stationary and exponential growth phases. (A) Kinetic of P*_hrtBA_* induction in stationary phase. HG001 WT and Δ*hrtBA* strains transformed with pP*_hrtBA_*-GFP. ON culture in CDM were distributed in a 96 well microplate. OD_600_ and GFP expression were followed in a spectrofluorimeter Infinite (Tecan). Results of hemin induced fluorescence minus non-induced (background, 0 µM hemin) are displayed. Results represent the average ± S.D. from triplicate biological samples. (B) Kinetic of HrtB expression following addition of hemin in HG001 WT. Strain from ON culture was diluted to OD_600_ = 0.1 in BHI. 2 µM hemin were added to the culture at t=0. At the indicated time points, samples of bacteria (containing equivalent number of bacteria) were pelleted and OD normalized to 0.5 by resuspending the pellet in PBS. At the indicated time point, samples of bacteria were pelleted and processed for SDS-PAGE and immunoblot with a α-HrtB antibody.

**Figure S2.**
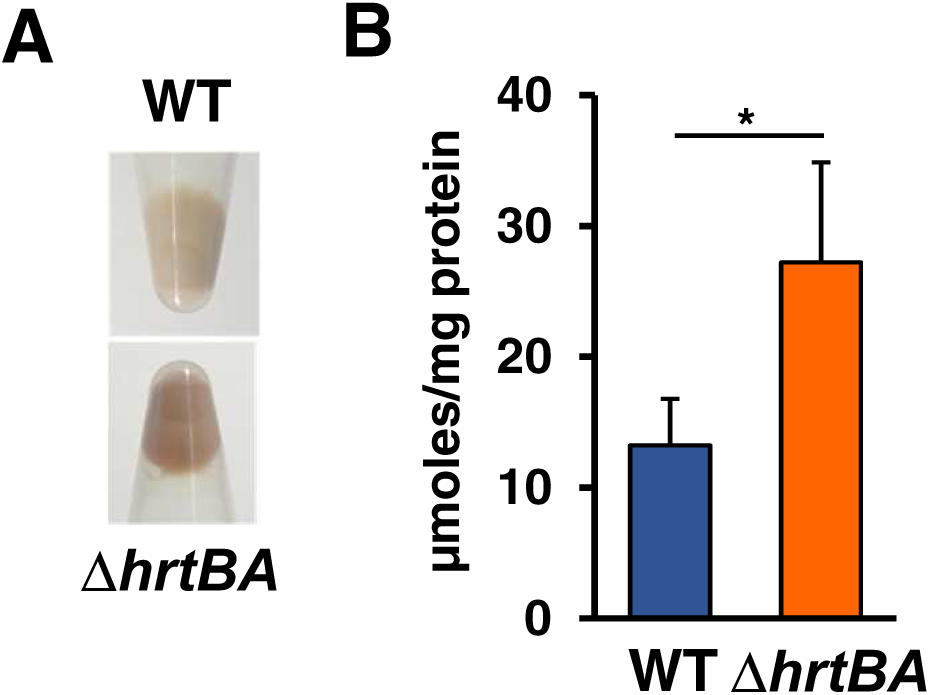
Heme accumulates in HG001 Δ*hrtBA* strain. WT and Δ*hrtBA* strains were grown to OD600 = 1 prior addition of 10 µM hemin in BHI for an additional 1.5 h. (A) Bacteria were pelleted by centrifugation and photographed. (B) Heme content in the cell pellets was determined by the pyridine hemochrome assay on lysates. Background from bacteria non-exposed to hemin was substracted. Results represent the average ± S.D. from biological triplicates. *, *P =0.045*, Student’s *t* test.

**Figure S3.**
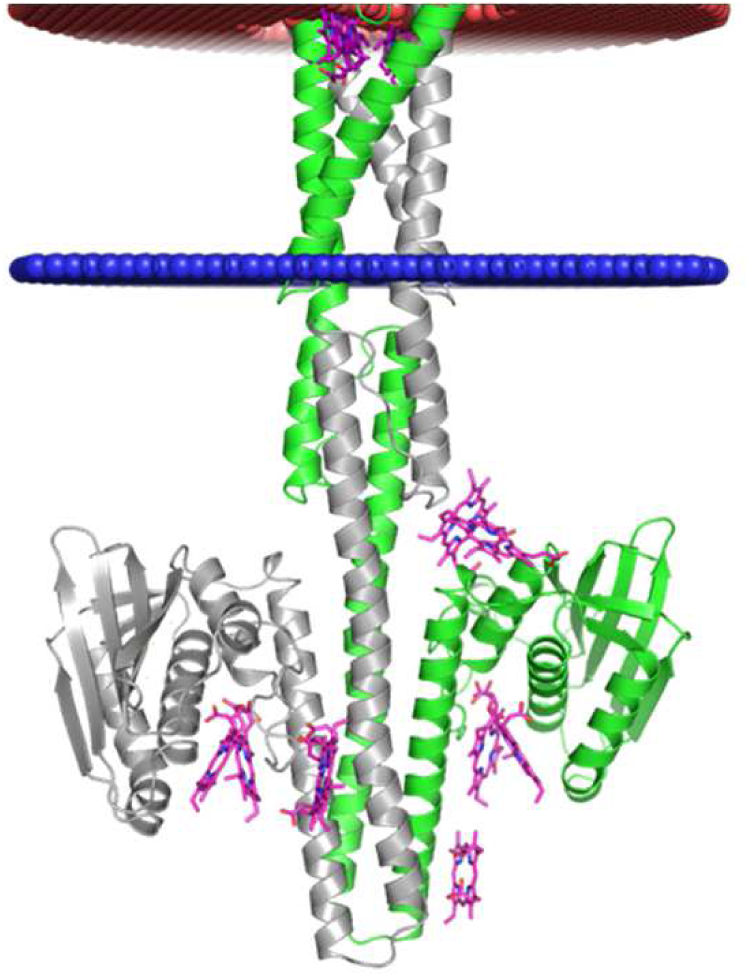
Superimposition of all the docking solutions using the intracellular part of HssS. As compared to Fig 3B using the ECD of HssS, all the docking solutions are above −8 kcal/mol and are scattered on the surface of the protein.

**Figure S4.**
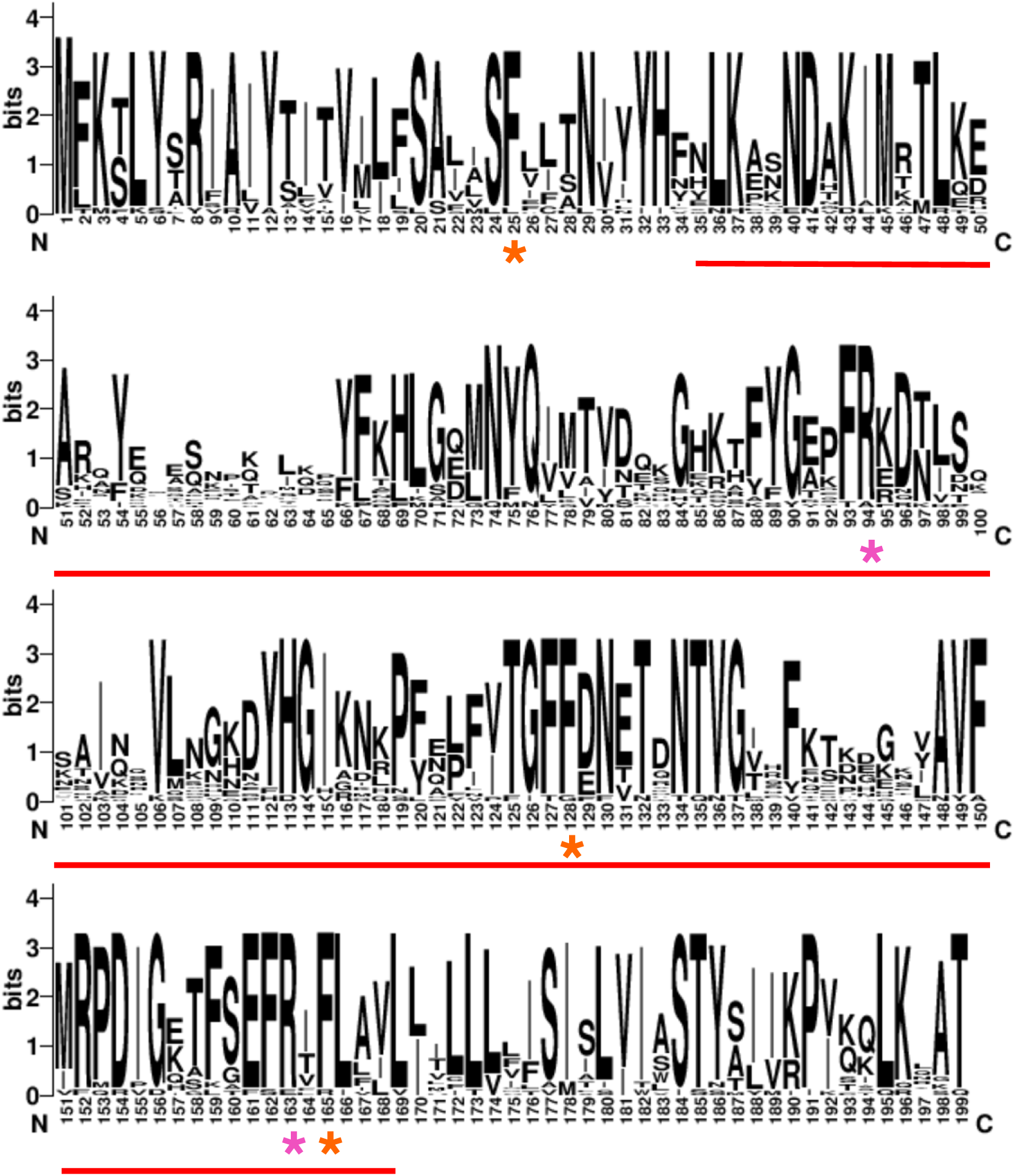
WebLogo representation of AAs [1-188] of HssS. Residues targeted by site-directed mutagenesis are highlighted by a red asterisk (orange for phenylalanine, pink for arginines). Red line represents the ECD.

**Figure S5.**
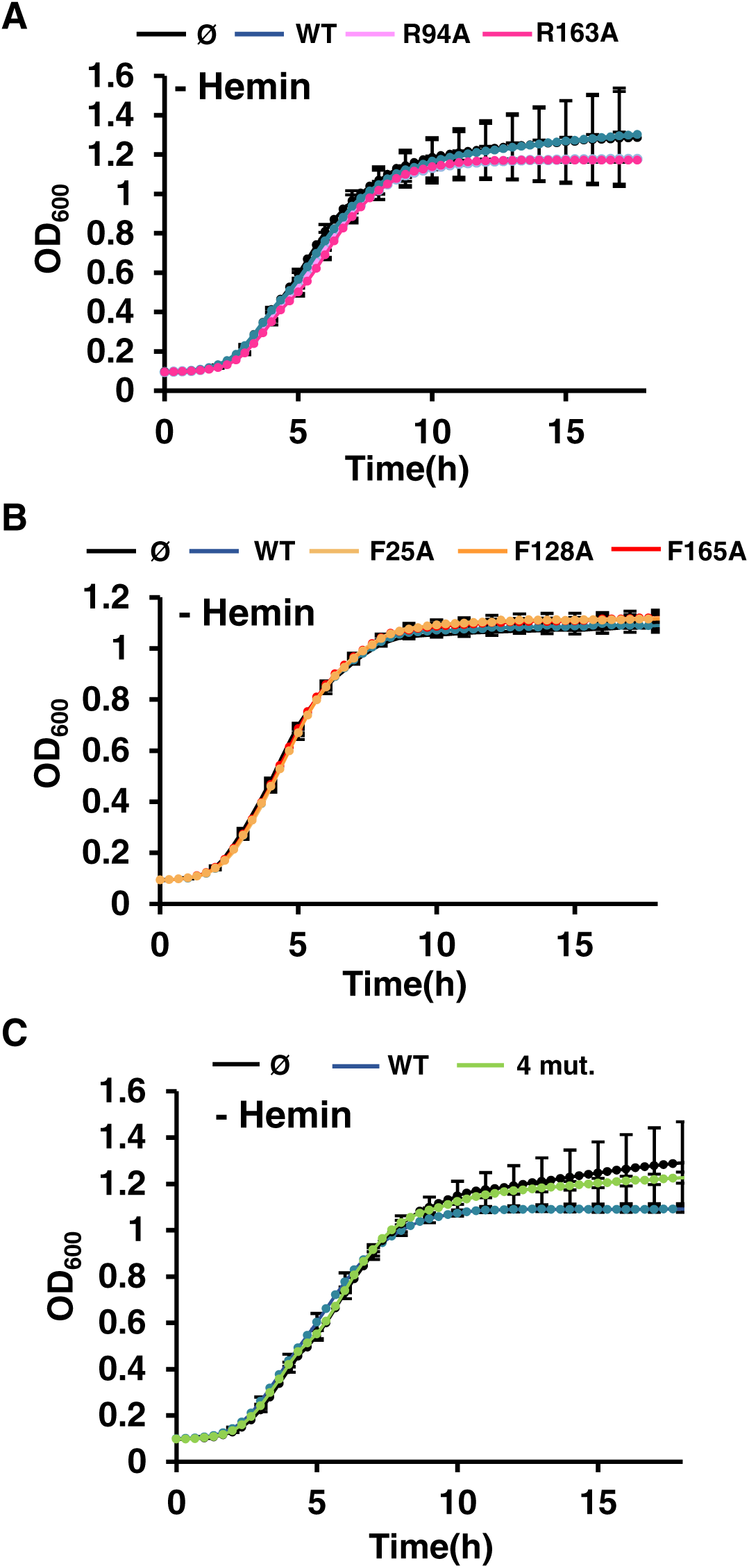
Growth of HG001 Δ*hssRS* complemented either with pGFP(HssS) or HssS variants R94A, R163A, F25A, F128A, F165A. All strains were were diluted from an ON preculture to an OD_600_ of 0.01 in CDM and grown in a microplate. OD_600_ was recorded every 20 min for the indicated time in a spectrophotometer (Spark, Tecan). (A) Growth of Δ*hssRS* mutants transformed either with pGFP(HssS), pGFP(HssS R94A), pGFP(HssS R163A), pGFP(HssS R94A, R163A) or empty vector (pØ). Results represent the average ± S.D from triplicate biological samples. (B) Growth of Δ*hssRS* mutants transformed either with pGFP(HssS), pGFP(HssS F25A), pGFP(HssS F128A), pGFP(HssS F165A) or empty vector (pØ). Results of bacterial growth minus medium background are displayed. Results represent the average ± S.D from triplicate biological samples. (C) Growth of Δ*hssRS* mutants transformed either with pGFP(HssS), pGFP(HssS 4 mut.) or empty vector (pØ). Results of bacterial growth minus medium background are displayed. Results represent the average ± S.D from triplicate biological samples.

**Figure S6.**
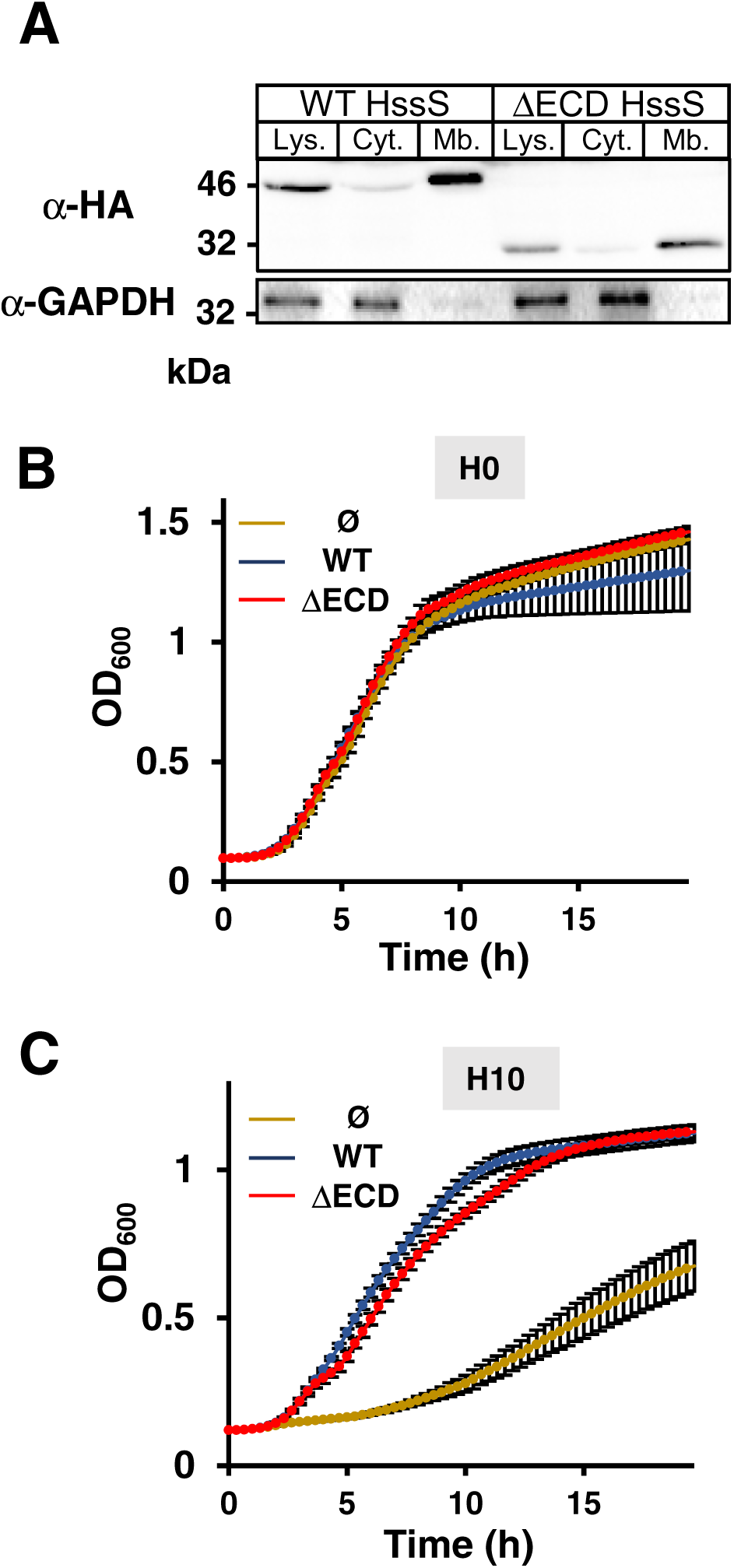
HssS ΔECD is expressed at the membrane and signals heme detoxification. (A) Cell fractionation of HssS FL and ΔECD expressing HG001 bacteria. Δ*hssRS* HG001 transformed with either pGFP(HssS) or pGFP(HssS ΔECD) were grown in BHI to an OD_600_ = 1, lysed and processed for the separation of membrane from cytoplasm by ultracentrifugation. Whole cell lysate, isolated membrane and cytoplasm enriched fractions were processed for SDS-PAGE and immunoblot using an anti-HA antibody. An anti-GAPDH antibody was used as a marker of cytoplasm. Result is representative of 3 independent experiments. (B-C) Heme toxicity in liquid culture of Δ*hssRS* HG001 transformed with either pGFP(HssS), pGFP(HssS ΔECD) or empty vector (pØ). Bacteria were diluted from ON cultures to an OD_600_ = 0.01 and grown in BHI without (B) or with 10 µM hemin in a microplate spectrophotometer (Spark, Tecan). OD_600_ was recorded every 15 min for the indicated time. Results of bacterial growth minus medium background are displayed. Results represent the average ± S.D from triplicate biological samples.

